# Transcriptional boosting and buffering during neurodevelopment by shifts in mRNA instability

**DOI:** 10.1101/2023.03.01.530249

**Authors:** Marat Mufteev, Deivid C. Rodrigues, Gwyneth Bolter, Kyoko E. Yuki, Ashrut Narula, Wei Wei, Alina Piekna, Jiajie Liu, Peter Pasceri, Olivia S. Rissland, Brett Trost, Michael D. Wilson, James Ellis

**Affiliations:** Program in Developmental Stem Cell, and Cancer Biology, Hospital for Sick Children, Toronto, Ontario M5G 0A4, Canada; Department of Molecular Genetics, University of Toronto, Toronto, Ontario M5S 1A8, Canada; Program in Molecular Medicine, Hospital for Sick Children, Toronto, Ontario M5G 0A4, Canada; Program in Genetics & Genome Biology, Hospital for Sick Children, Toronto, Ontario M5G 0A4, Canada; RNA Bioscience Initiative and Department of Biochemistry & Molecular Genetics, University of Colorado School of Medicine, Aurora, Colorado 80045, USA

**Author notes:** equal contribution (Marat Mufteev, Deivid C. Rodrigues, Gwyneth Bolter).

**Keywords:** transcription rate, RNA half-life, neurodevelopment, microRNA, RNA-binding protein, transcriptional buffering, transcriptional boosting, post-transcriptional regulation, 3ʹUTR length

## Abstract

The contribution of mRNA half-life is commonly overlooked when examining changes in mRNA abundance during development. mRNA levels are regulated by transcription rate or mRNA half-life, but combinations of both could boost or buffer the final steady-state. We measured transcription rate and mRNA half-life changes during human induced pluripotent stem cell (iPSC)-derived neuronal development using RATE-seq. The most prevalent mode of gene regulation during transitions to neuronal progenitors and neurons was transcriptional buffering. Less prevalent modes consisted of transcriptional boosting events, transcription rate only shifts, or mRNA half-life only shifts. Buffered and novel boosted gene sets were enriched in RNA-binding protein (RBP) motifs and microRNA sites. Global mRNA half-life decreased two-fold in neurons. microRNA copy number per cell increased during neurodevelopment coincident with destabilization of a subset of short isoforms that contributed to increased average 3ʹUTR length. Our findings identify mRNA instability mechanisms that regulate transcript levels and 3ʹUTR isoform abundance, and provide a precedent for transcriptional boosting or buffering during development of other human tissues.

## INTRODUCTION

Regulation of mRNA stability is essential for pluripotent stem cell homeostasis (Jia *et al*., 2012; Li *et al*., 2015) and neuronal development (Prashad and Gopal, 2021). mRNA stability also regulates many genes that impact cortical neuron activity, including autism spectrum disorder-risk genes (Lee *et al*., 2016) and Fragile X Messenger Ribonucleoprotein 1 (FMRP) targets (Shu *et al*., 2020). During development, active modulation of mRNA stability can fine-tune transcriptional programs to make them more precise (Ebert and Sharp, 2012). Threshold effects evoked by changes in microRNA abundance relative to their target mRNA levels is one such mechanism (Mukherji *et al*., 2011; Ebert and Sharp, 2012; Bosson *et al*., 2014). Many RNA binding proteins (RBPs) participate in regulating mRNA stability, with examples such as ELAVL1 being able to dissociate microRNA-Target interactions or influence alternative polyadenylation site choice (Kundu *et al*., 2012; Wei *et al*., 2020). More broadly, coordination between mRNA transcription and stability has been shown to buffer transcript abundance from yeast to human cells in response to environmental or genetic manipulations (Lee *et al*., 2012; Timmers and Tora, 2018). Such transcriptional buffering events occur when changes in transcription rates are compensated for by opposing changes in mRNA half-life. Transcriptional boosting events involve same-direction changes in transcription rate and mRNA half-life, but have not been reported in human disease or investigated during human development.

To understand the dynamics of transcriptome regulation as healthy cells differentiate through developmental stages, it is crucial to characterize transcription rates and mRNA stability on a genome-wide scale. To search for evidence of changes in mRNA half-life that can buffer or boost changes in transcription rates during human neurodevelopmental stages, pluripotent stem cells can be differentiated *in vitro* into neural progenitor cells (NPC) and cortical neurons (Brennand *et al*., 2011). We previously used induced Pluripotent Stem Cells (iPSC) to produce cortical neurons from an individual with Rett syndrome (RTT) compared to their isogenic control and demonstrated pronounced mRNA half-life only shifts and transcriptional buffering of hundreds of genes in this neurodevelopmental disorder (Rodrigues *et al*., 2023). We found that in RTT neurons, certain RBP-binding motifs were enriched in buffered gene sets where the transcription rate was up and the half-life was down. Buffering has also been found in mouse embryonic stem cells (ESC) that are deficient for subunits of transcriptional co-activator complexes (Forouzanfar *et al*., 2025). Transcriptional boosting was not observed in these studies.

Powerful techniques for genome-wide measurement of mRNA half-life at high resolution use metabolic labeling of RNAs followed by sequencing. These methods include thiol (SH)-linked alkylation for the metabolic sequencing of RNA (SLAM-seq) (Herzog *et al*., 2017) and RNA approach to equilibrium sequencing (RATE-seq) (Neymotin *et al*., 2014). RNA half-life is then measured from decay or accumulation of mRNAs following label incorporation of up to 24 hours or pulse-and-chase of modified RNA nucleotides. In addition, RNA half-life can be deduced from the equilibrium relationship between steady-state abundance and transcription rate. In this case, transcription rate is measured using samples from the short labeling times before RNA degradation (Muhar *et al*., 2018). Moreover, 3ʹ-end sequencing of metabolic labeled RNAs accurately defines the length of each 3ʹUTR isoform of a gene allowing analysis of mRNA half-life at the isoform level. Genome-wide mRNA half-life measurements have been performed across different cell types and organisms. However, most studies measured mRNA half-life in a single cell type or compared perturbed cell conditions (Schwalb *et al*., 2016; Muhar *et al*., 2018; Schofield *et al*., 2018). The developmental dynamics of transcription rate and mRNA half-life shifts during human neurodevelopment that contribute to transcriptional buffering, boosting and 3ʹUTR isoform length are unknown.

Most eukaryotic genes contain multiple 3ʹ-end cleavage and polyadenylation sites. It is well-established in several organisms that neurogenesis is accompanied by the utilization of longer 3ʹUTR isoforms (Hilgers *et al*., 2011; Smibert *et al*., 2012; Ulitsky *et al*., 2012). Although unusually long 3ʹUTR isoforms are found in neurons (Miura *et al*., 2013), single-cell RNA-seq analysis during mouse embryogenesis revealed a gradual increase in 3ʹUTR length from E9.5 to E13.5 in most tissues (Agarwal *et al*., 2021). A functional role of 3ʹUTR length regulation during development is supported by evidence that knockdown of the mRNA cleavage machinery component *NUDT21* increases the reprogramming efficiency of human fibroblasts into iPSC (Brumbaugh *et al*., 2018).

Differences in the stability of 3ʹUTR isoforms can also control their relative abundance, as is the case with the critical neuronal transcription modulator *MECP2*. In human ESC (hESC), the RBP PUM1 and pluripotent-specific microRNAs actively degrade *MECP2* mRNAs containing the long 3ʹUTR. This degradation halts in neurons leading to an increase in abundance of the long 3ʹUTR isoform of *MECP2* (Rodrigues *et al*., 2016). This observation suggests that 3ʹUTR isoform length can be determined by shifts in mRNA stability, and this can be evaluated using 3ʹ-end sequencing. Other potential mechanisms could act to make short isoforms, which have fewer RNA motifs, more sensitive to degradation by increasing the levels of cell-specific motif-binding molecules such as microRNA beyond a threshold (Ebert and Sharp, 2012).

Here, we systematically measured mRNA transcription rate and half-life at the 3ʹUTR isoform level using RATE-seq at three stages of human neurodevelopment: iPSC, NPC and cortical neurons. We revealed a widespread influence of mRNA stability shifts on the landscape of mRNA isoforms initially set by transcription rates in iPSC and during the early stages of human neuronal development. Transcriptional buffering by compensatory half-life changes was observed in up to 50% of genes with changed transcription rates, and transcriptional boosting by additive half-life changes occurs in up to 15% of genes. RBP and microRNA binding motifs were enriched on buffered and boosted genes. Half-life only changes had roughly the same frequency as transcription rate only changes. Overall, 80% of altered genes exhibited mRNA stability changes during neurodevelopment, and 50% of genes targeted by pluripotent transcription factors in iPSC were buffered or boosted by half-life changes. We also report reduced global transcript half-life in neurons, an increased microRNA abundance per cell and destabilization of a subset of short isoforms in neurons relative to pluripotent cells, coincident with the apparent lengthening of 3ʹUTR isoforms during human neurodevelopment.

## RESULTS

### RATE-seq captures global shifts in transcription rate and RNA steady-state during human neurodevelopment

To directly determine the contribution of transcription rates and RNA stability on mRNA steady-state abundance and 3ʹUTR isoform usage, we performed RATE-seq using a 3ʹ-end sequencing method (QuantSeq) on cells collected and processed at different stages of iPSC-based neuronal development. The original RATE-seq approach measures absolute mRNA half-life from the incorporation kinetics of the modified nucleotide 4sU into newly transcribed nascent mRNAs over time points of up to 24 hours (Neymotin *et al*., 2014). We have recently adapted RATE-seq to estimate a relative mRNA half-life by calculating the fold-change ratio between steady-state and transcription rate measured from early time points of 0.5 and 1 hour (Rodrigues *et al*., 2023) (Fig 1A).

**Figure 1.**
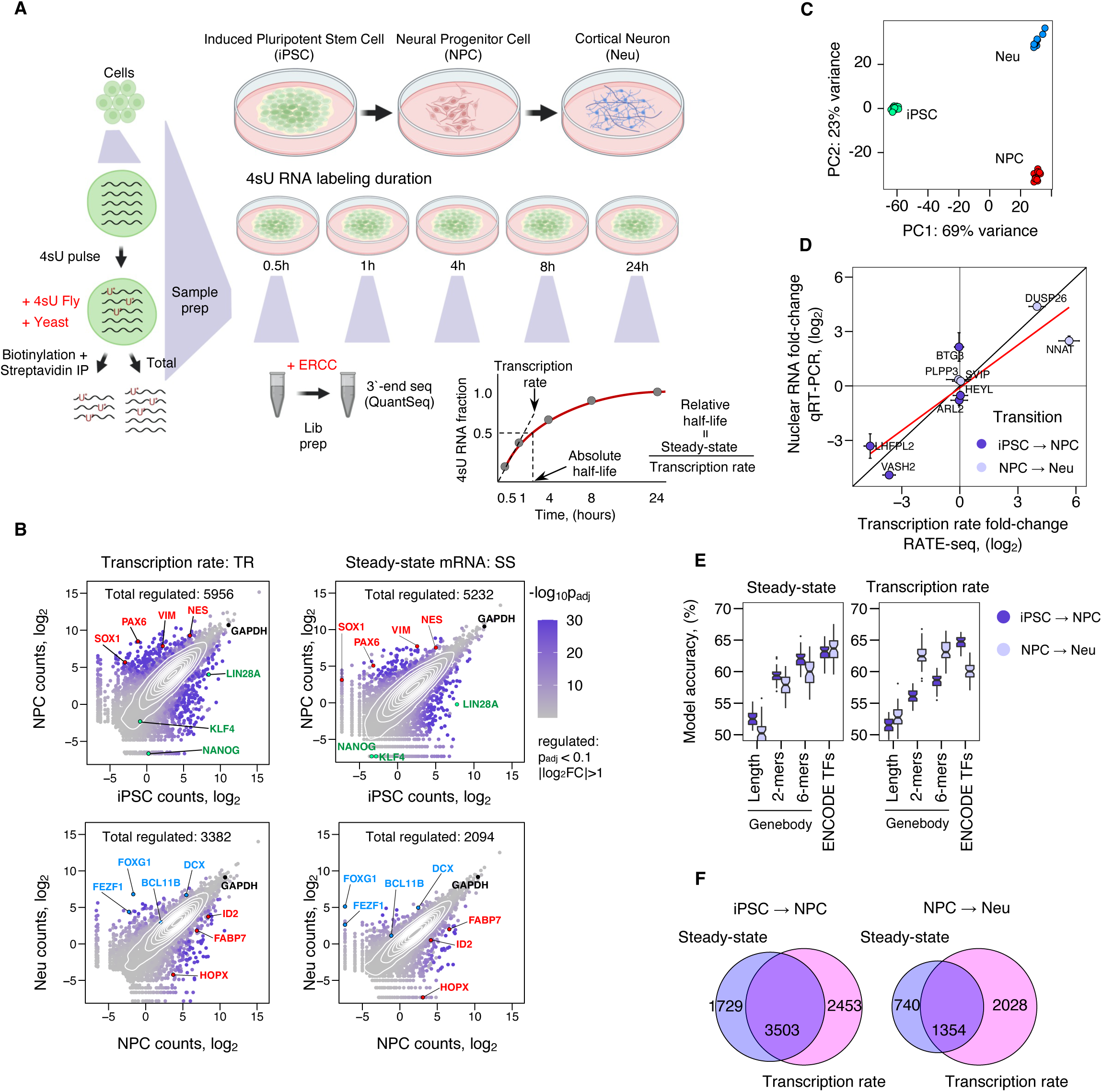
Changes in transcription rate during neurodevelopment do not always result in altered mRNA steady states. A, schematics of experimental outline (created with BioRender.com) for simultaneous quantification of transcription rate, mRNA half-life, and steady-state mRNA level. iPSC and iPSC-derived NPC and cortical neurons were pulse-labeled with 4sU, and at designated time points total RNA was harvested. 4sU-labeled *Drosophila melanogaster* (fly) and unlabeled *Saccharomyces cerevisiae* (yeast), and ERCC spike-in RNAs were added as indicated and used as pull-down efficiency, non-specific binding, and library preparation and sequencing controls, respectively. Steady-state mRNA levels were quantified from an aliquot of the 24-hour time point (non-biotinylated and unprocessed). Two replicates of this experiment were performed. B, scatterplots depicting genome-wide changes in transcription rate and steady-state after both iPSC-to-NPC and NPC-to-Neu transitions. Cell type marker genes for iPSC, NPC, Neu are highlighted by green, red, and blue colours, respectively. C, principal component analysis of sequencing data from iPSC, NPC, and Neu showing significant separation between cell types. D, transcription rate fold-changes determined by RATE-seq (x-axis) were validated using an alternative approach. Nuclear RNA fold-change measured by qRT-PCR (y-axis) was used as a proxy for transcription rate shifts of genes selected to cover a large spectrum of fold-changes, including genes with no changes (n = 2). E, random forest classifier for predicting steady-state and transcription rate direction based on gene-body sequence and promoter occupancy by ENCODE transcription factors (TF). Each point represents a classifier’s accuracy when trained on a particular sample of training and test data. A boxplot for a model is obtained from 50 resamplings. F, overlap of genes altered at transcription rate and/or steady-state during neuronal development.

Cells were collected and processed simultaneously at three stages of human neurodevelopment *in vitr*o. We used a control iPSC line (d3-4 #37) and NPC derived from them using a dual SMAD differentiation protocol. Cortical neurons (Neu) were matured from the NPC for 4 weeks. The neurons were magnetically sorted based on the presence of specific cell surface proteins to exclude contaminant glia and progenitor cells (Rodrigues *et al*., 2020). RATE-seq data from these same control neuron samples have already been reported in comparison to isogenic RTT neurons (Rodrigues *et al*., 2023). Total RNA from all three cell types was collected at 5 different time points (0.5, 1, 4, 8 and 24 hours) in two replicate experiments, biotinylated and pulled down using streptavidin beads (Neymotin *et al*., 2014). Exogenous 4sU labeled fly and unlabeled yeast whole-cell RNAs were spiked into the samples to reconstruct the human 4sU labeled fraction within the total mRNA pool and to control for pull-down efficiency and background contamination, respectively (Fig S1A-C). External RNA controls consortium (ERCC) spike-in controls were added during the 3ʹ-end sequencing library preparations. Transcription rate was measured at 0.5-hour and 1-hour timepoints when mRNA degradation has a negligible effect on the labeled RNAs for most genes (Muhar *et al*., 2018). In addition, steady-state abundance was measured from the input sample obtained at the 24-hour time point, matching the 4sU incubation conditions between samples. As expected, incubation with 4sU for 24 hours did not affect the viability of cells (Fig S1D).

We found approximately 2100-6000 genes with changes in transcription rate or steady-state, or both, after neurodevelopmental transitions of iPSC-to-NPC and NPC-to-Neu (Fig 1B) (Supplementary table 1). Supporting the ability of the differentiation protocol to produce NPC and neurons, we observed a decrease in transcription rate in pluripotency marker genes and an increase in progenitor marker genes in the first transition, and then a decrease in progenitor marker genes and an increase in neuron marker genes in the second transition (Fig 1B). Similarly, mRNA abundance of progenitor and then neuron marker genes increased respectively at the steady-state upon the transitions while housekeeping genes were unchanged (Fig 1B). Principal component analysis (PCA) separated cell types into distinct clusters, and the abundance of endogenous and ERCC spike-in RNAs was highly reproducible between replicates (Fig 1C and Fig S1E-F). Transcription rate shifts were validated for a selection of genes by qRT-PCR on nuclear fractions collected from independently differentiated cells (Fig 1D). Finally, we checked whether gene-body length, k-mers (dinucleotides or 6-mers) or promoter occupancy by transcription factors (ENCODE TFs (ENCODE Project Consortium, 2011)) obtained from Harmonizome (Ben-Porath *et al*., 2008) can predict the direction of the transcription rate and steady-state shifts. We used a sequence content-based random forest classifier to predict the effect of these features on steady-state abundance and transcription rate. No predictive ability above random chance was observed for gene-body length, but an approximately 60% accuracy was found for 6-mers and the ENCODE TFs on the steady-state during both transitions (Fig 1E). The best predictor of changes in the direction of transcription rate during the iPSC to NPC transition was ENCODE TFs at 65% accuracy, but 6-mers were more accurate than ENCODE TFs in the NPC-to-Neu transition.

Importantly, change in transcription rate for 2453 and 2028 (approximately 40%-60% of all genes with transcription rate changes in Fig 1F) led to no significant change in their steady-state levels in both cell transitions. This result emphasizes that about half of transcription rate changes led to altered steady-state abundance, while the other half were buffered at the steady-state level. Similarly, 1729 and 740 genes, respectively, had altered steady-state abundance but no corresponding shift in transcription rate, suggesting that these genes were differentially regulated at the level of mRNA stability (Fig 1F). Together, the observed discrepancy between transcription rate and steady-state shifts suggests a widespread contribution of mRNA stability in the regulation of genes during human neurodevelopment.

### Absolute and relative mRNA half-life changes identify buffered and novel boosted gene sets

Our profiling of transcription rate and steady-state changes indicated a genome-wide effect of mRNA stability during both neurodevelopmental transitions. To quantify this effect, we used the 4sU saturation kinetics over the entire 24-hour time course (Fig S1A). By measuring the absolute change in mRNA half-life in hours, we observed a mean mRNA half-life of 5.4 hours in iPSC and NPC, compared to 2.9 hours in neurons (Rodrigues *et al*., 2023) (Fig 2A).

**Figure 2.**
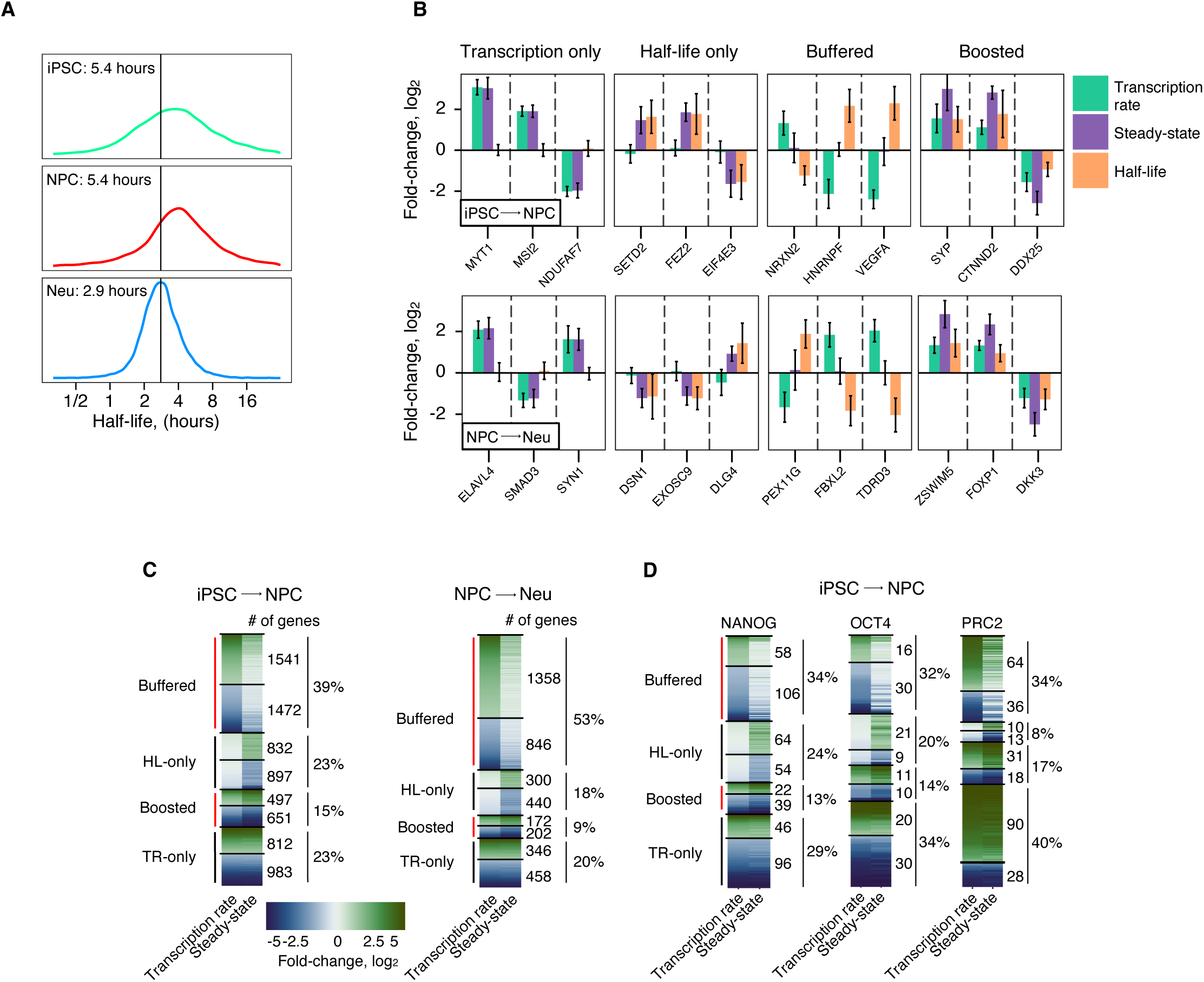
Widespread changes in mRNA half-life directly alter steady-state abundance to buffer or boost transcription rate changes during neuronal development. A, global decrease in the absolute mRNA half-life after transition to Neu. The mean half-life for each cell type is shown in the figure panel. B, examples of genes with changes in transcription rate or half-life only, or buffered or boosted by half-life changes. C, number and percentage of all genes with changes in the 4 modes of regulation during cell transitions: buffered, half-life (HL) only, boosted, and transcription rate (TR) only. The height of each group corresponds to the number of genes. D, number and percentage of genes targeted by the pluripotent TF denoted on top with changes in the 4 modes of regulation during the iPSC-to-NPC transition.

We also used steady-state and transcription rate ratios to derive the relative shifts in mRNA half-life during neurodevelopment. The ratio and saturation curve methods employed to measure mRNA half-life were concordant with each other for the iPSC and NPC, similar to our findings with the Neu samples (Rodrigues *et al*., 2023) (Fig S2A-C, Supplementary Table 2). However, the absolute method (mRNA half-life in hours) required a sufficiently high transcription rate to measure the saturation curves accurately and thus yielded fewer quantified genes (Fig S2A). This was in contrast to the ability of the ratio method to measure relative fold-changes in genes with low transcription rates, as already shown in the neuron samples (Rodrigues *et al*., 2023). We observed that shifts in mRNA half-life during both cell transitions resulted in four modes of gene regulation: 1) steady-state changes modulated by transcription rate only; 2) steady-state changes caused by mRNA half-life only; 3) transcription rate changes buffered by reciprocal changes in mRNA half-life; or 4) transcription rate changes boosted by same-direction changes in mRNA half-life (Fig 2B-C).

We found that for approximately half of the regulated genes the mRNA half-life compensated for shifts in transcription rate resulting in buffering of steady-state levels after both developmental transitions. We observed 3013 genes (39%) after iPSC-to-NPC and 2204 genes (53%) after NPC-to-Neu transitions were buffered where mRNA stability shifts counteracted the changes in transcription (Fig 2C). Interestingly, another 1729 (23%) and 740 (18%) genes regulated steady-state through half-life only mechanisms after each developmental transition (Fig 2C). A further 1148 (15%) and 374 (9%) genes boosted transcription rate shifts through mRNA half-life, leading to amplified steady-state changes (Fig 2C). Finally, we detected 1795 (23%) and 804 (20%) genes with changes in steady-state mediated by transcription rate only changes.

Since ENCODE TFs partially predicted the direction of transcription rate and steady-state changes in the iPSC-to-NPC transition (Fig 1E), we examined whether networks of genes targeted by core pluripotency TFs escaped buffering and boosting. We first obtained gene targets of *NANOG*, *OCT4*, and Polycomb Repressive Complex 2 (PRC2) which were defined as overlapping SUZ12, EED, H3K27 targets reported in previously published ChIP-seq experiments in hESC (Ben-Porath *et al*., 2008). Approximately 30-40% of target genes bound by core pluripotency TFs were regulated by transcription rate only changes as iPSC differentiate into NPC (Fig 2D). In support of our transcription rate data, we found that the PRC2 targets were predominantly upregulated in NPC, supporting the repressive role of these TFs in iPSC (Fig 2D). As observed above with the 4 global modes of gene regulation, mRNA half-life changes still buffered or boosted transcription rate changes in nearly 50% of genes that are TF targets of NANOG, OCT4, or PRC2 (Fig 2D). The remaining 20% of NANOG and OCT4 target genes were regulated by mRNA half-life only. PRC2 had a >2-fold reduction in target genes controlled by mRNA half-life only (down to 8%). Together, our data highlight the widespread contribution of half-life in buffering and boosting mRNA levels (as observed in Fig 1E) as stem cells differentiate into progenitors and neurons. Buffering caused by shifts in half-life in different cell types leads to a substantial decoupling of steady-state levels from the transcription rate changes of these genes.

### Validation of buffering and boosting in subcellular fractionation datasets

To orthogonally validate our findings of buffered and boosted gene sets, we reanalyzed published RNASeq datasets derived from hESC, NPC and neurons after nuclear and cytoplasmic subcellular fractionation (Blair *et al*., 2017). This approach uses the nuclear fraction as a proxy for transcription rate and the cytoplasmic fraction as a proxy for steady-state mRNA levels, allowing for the calculation of relative half-life fold changes (Fig 3A). We previously demonstrated this method when validating transcriptional buffering in mouse models of RTT (Rodrigues *et al*., 2023). In the first transition from hESC-to-NPC, genes in the buffered or boosted mode had similar percentage values (Fig 3B, Supplementary table 3). Reassuringly, despite a different pluripotent stem cell line, differentiation methods, and method of measuring half-life that does not employ 4sU, comparisons of all genes with our RATEseq iPSC-to-NPC datasets show statistically significant overlaps for each of the 4 modes of regulation. Similarly, in the NPC-to-Neu transition, we observed significant overlaps for each mode of regulation with our RATEseq data (Fig 3C, Supplementary table 3). While the Blair et al (2017) dataset included both 14-day and 50-day-old neurons, both of which exhibited all 4 modes of regulation (Fig S3), here we focused on the 14-day neurons as their developmental stage more closely aligns with our 4-week iPSC-derived neurons. These findings confirm that boosting and buffering are reproducible features of human neurodevelopment.

**Figure 3.**
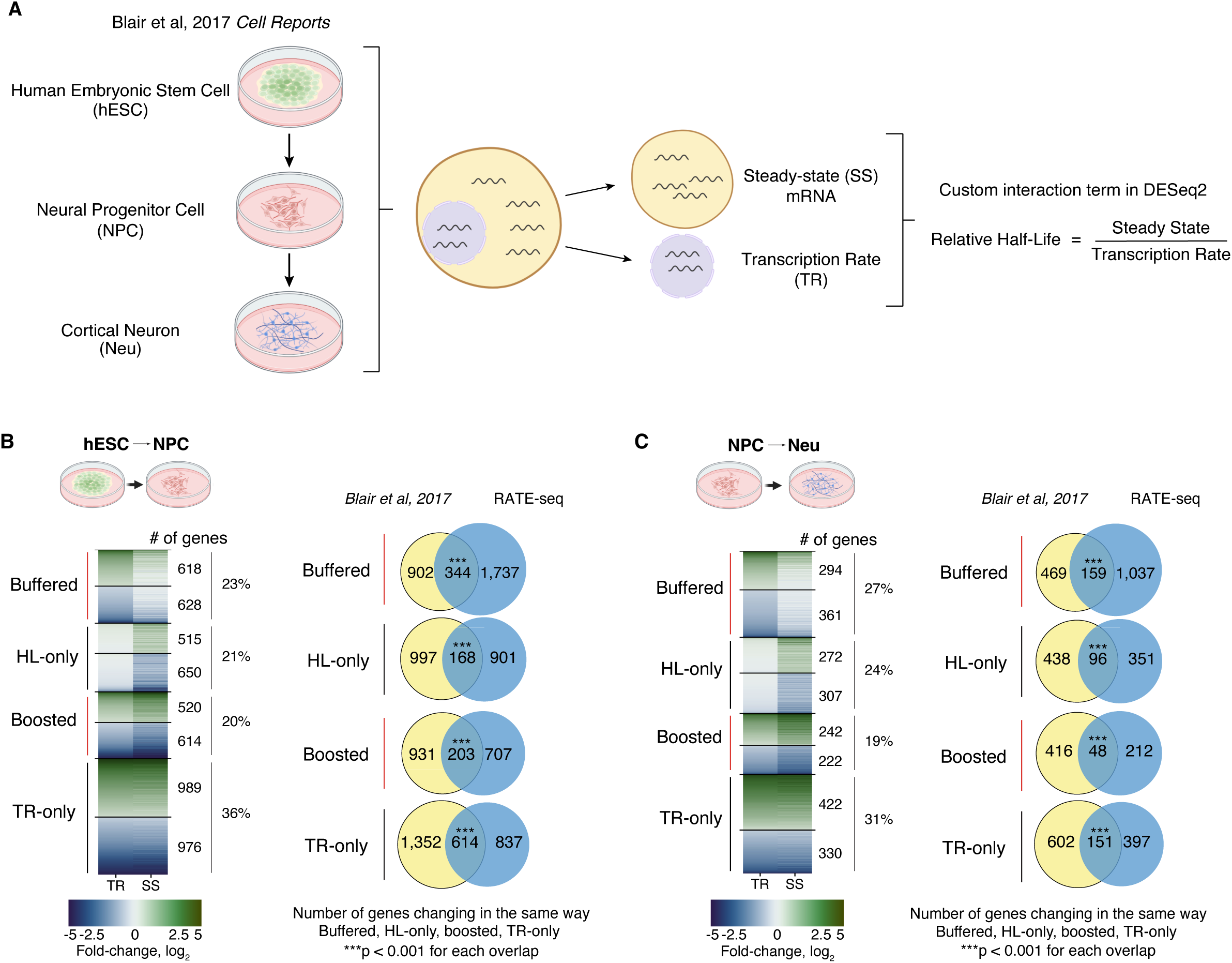
Buffered and boosted genes are observed in neurodevelopment subcellular fractionation datasets. A, Rationale behind subcellular fraction usage for mRNA transcription rate and half-life measurements after hESC differentiate into NPC and cortical neurons (D14) (Blair *et al*., 2017). RNAseq was analyzed using a custom interaction term in DESeq2 to compare ratios between the fractions. B, Left panel - number and percentage of all genes with changes in the 4 modes of regulation during the hESC-to-NPC cell transition: buffered, half-life (HL) only, boosted and transcription rate (TR) only. The height of each group corresponds to the number of genes. Right panel - fraction of genes exhibiting the same type of change in RATE-seq (this study) and Blair et al 2017 (right) for each of the 4 modes of regulation. Overlaps were calculated using a hypergeometric test. C, Left panel - number and percentage of all genes with changes in the 4 modes of regulation during the NPC-to-Neu cell transition: buffered, HL-only, boosted and TR-only. The height of each group corresponds to the number of genes. Right panel - fraction of genes exhibiting the same type of change in RATE-seq (this study) and Blair et al 2017 (right) for each of the 4 modes of regulation. Overlaps were calculated using a hypergeometric test.

### RBP motifs are enriched in buffered and boosted genes

Our previous analysis demonstrated that mRNA motifs for RBPs capable of shuttling between the nucleus and cytoplasm are overrepresented in genes undergoing transcriptional buffering in RTT neurons (Rodrigues *et al*., 2023). These RBP motifs were enriched on buffered genes with half-life (HL) down in RTT neurons, and no motifs were discovered for buffered genes with HL up. We performed the same RBP motif enrichment analysis for the 174 RBP *cis*-acting elements on the buffered genes in the new neurodevelopmental RATE-seq data. In the iPSC-to-NPC transition, the buffered genes with HL down were enriched for nuclear-cytoplasmic shuttling RBP motifs that we previously reported for RTT neurons (Fig 4A, Supplementary table 4). However, there were also new RBP motifs now enriched on the buffered HL up genes, which included ELAVL, IGF2BP, and PABPC motifs (Fig 4A). In contrast, in the buffered genes with HL down, these motifs were depleted and LIN28, SAMD4, PPRC1, NCL and YBX motifs were enriched instead. In iPSC-to-Neu, the RBP motifs in the buffered genes resembled those found in the iPSC-to-NPC transition (Fig 4A, Supplementary table 4). This finding suggests that NPC and neurons use similar RBP motifs for buffering, explaining why we found so few motifs enriched in the NPC-to-Neu transition (Fig S4). During neurodevelopment, RBP motifs were implicated in both HL up and HL down buffered gene sets.

**Figure 4.**
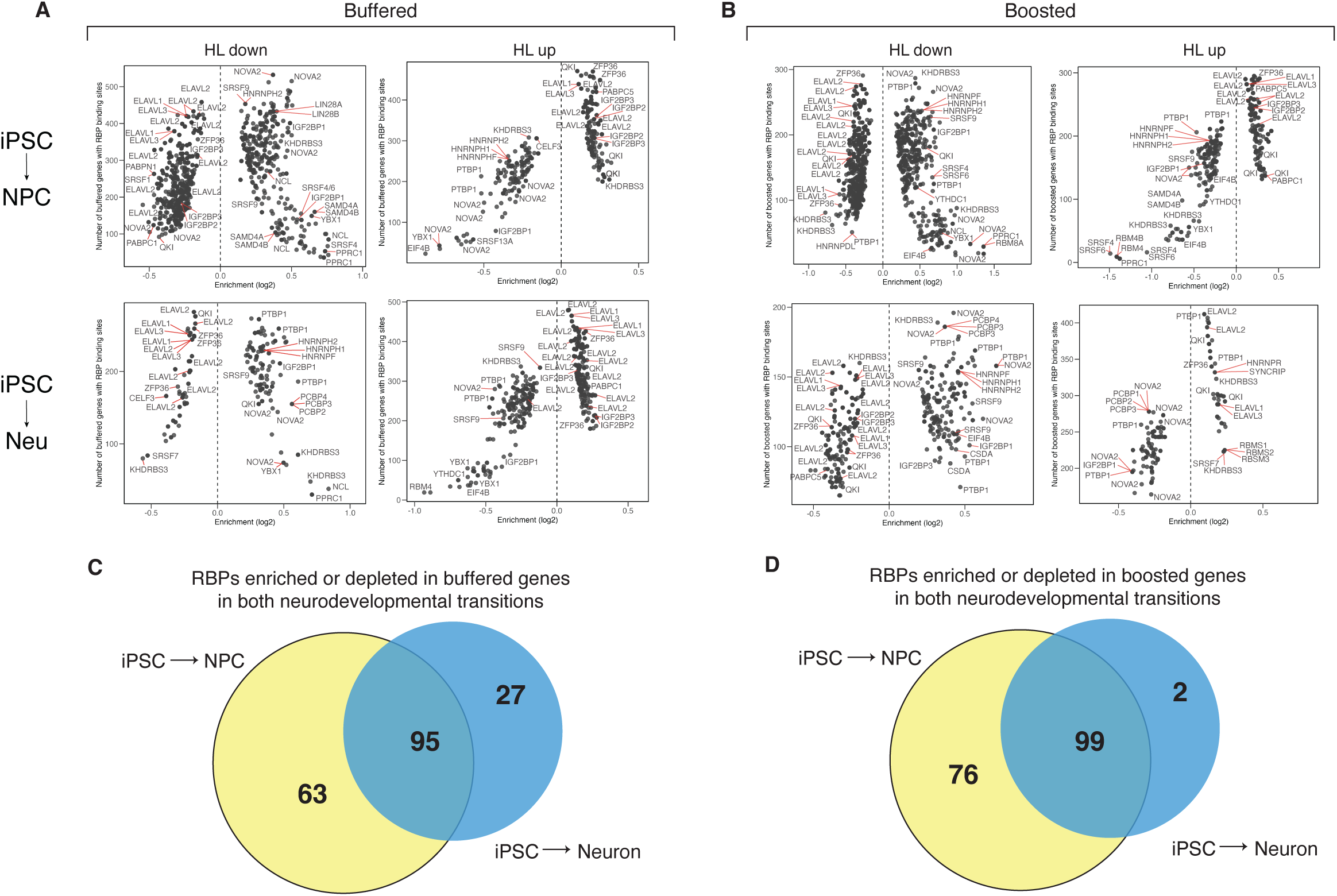
RBP motifs are enriched on buffered and boosted genes during neurodevelopment. A, 6-mers known to be targeted by RBPs enriched in the groups of buffered genes in iPSC-to-NPC (top) and iPSC-to-Neu (bottom). B, 6-mers known to be targeted by RBPs enriched in the groups of boosted genes in iPSC-to-NPC (top) and iPSC-to-Neu (bottom). C, RBP motifs enriched or depleted on buffered genes in iPSC-to-NPC (buffered HL up AND buffered HL down) that overlap with the enriched or depleted RBP motifs in iPSC-to-Neu. D, RBP motifs enriched or depleted in boosted genes in iPSC-to-NPC (boosted HL up AND boosted HL down) that overlap the enriched or depleted RBP motifs in iPSC-to-Neu.

We also performed the RBP motif enrichment analyses on the novel boosted genes. We detected the same RBP motifs that were found for the buffered genes, with boosted HL up resembling buffered HL up, and boosted HL down being similar to buffered HL down (Fig 4B, Supplementary table 5). There were similar enrichment or depletion plots of RBP motifs in the iPSC-to-NPC transition and in iPSC-to-Neu (Fig 4B, Supplementary table 5). Few enriched RBP motifs were found in the NPC-to-Neu transition (Fig S4). Examination of RBP motifs enriched or depleted in both iPSC-to-NPC and iPSC-to-Neu showed an overlap of roughly half for buffered and boosted genes (Fig 4C-D), supporting a role in both cell types. Overall, we describe the new RBP motifs associated with buffered HL up and novel boosted genes in human neurodevelopment, and extend previously reported RBP motifs for buffered HL down genes from disease to neurodevelopment.

### RNA instability contributes to 3ʹUTR lengthening during neurodevelopment

To define the contribution of mRNA instability to the specification of the 3ʹUTR usage during human neurodevelopment, we quantified mRNA transcription rate and half-life for all known 3ʹ-end mRNA isoforms. In this analysis, the transcription rate of a 3ʹUTR isoform is the net result of gene transcription and cleavage frequency at the corresponding pA site, measured in the first time points of 4sU labeling (0.5h and 1h). In addition, the high-resolution annotation defined by the roughly 200,000 pA sites previously compiled by Wang *et al* (Wang *et al*., 2018) allowed the comprehensive comparison between 3ʹUTR isoforms of the same gene. In particular, we detected that approximately 10,000 genes contained multiple 3ʹUTR isoforms in their last exon. For most genes, two 3ʹUTR isoforms with large distances of up to 4kb between the pA sites accounted for at least 80% of the total mRNA abundance (Fig S5A-B). Based on this observation, we focused the analysis on the 2 most abundant 3ʹUTR isoforms that differ by >500 nucleotides (nt) for each gene.

We define short or long 3ʹUTR-containing mRNAs as those cleaved and polyadenylated at the most proximal or distal sites to the stop codon, respectively (Fig 5A). We observed that, on average, processed short and long 3ʹUTR-containing mRNAs had equal transcription rates in iPSC. However, the long 3ʹUTR-containing mRNA isoforms were generally less abundant than short isoforms at the steady-state (Fig 5A-B and Fig S5C). These data suggest that active degradation of mRNAs with long 3ʹUTRs in iPSC is widespread, extending the published observation of the *MECP2* long 3ʹUTR-containing mRNA degradation in hESC (Fig S5C).

**Figure 5.**
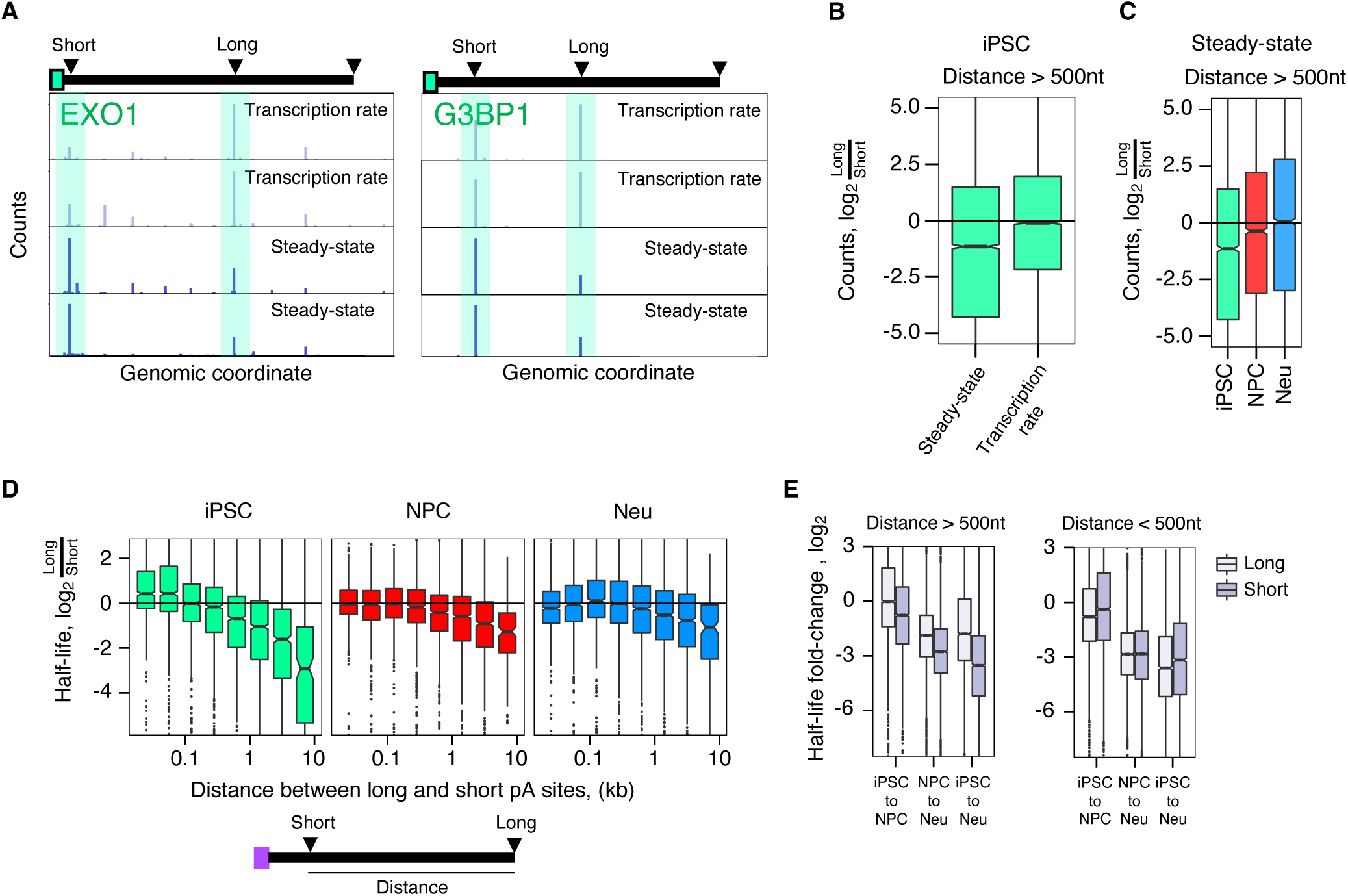
RNA instability affects the 3ʹUTR landscape in pluripotent stem cells and during neurodevelopment. A, representative sequencing read peaks in the iPSC transcription rate (upper) and steady-state (lower) samples, corresponding to polyadenylation sites (arrowheads) in the 3ʹUTR as an example of genes with active degradation of long 3ʹUTR-containing mRNAs. y-axis represents the number of sequencing reads (n = 2). B-C, comparison in detected abundance of long and short 3ʹUTR-containing mRNAs. Genes with a distance >500 nucleotides between polyadenylation sites corresponding to long and short 3ʹUTR-containing mRNAs are included in the boxplots. B, steady-state and transcription rate read counts fold-change (y-axis) between long and short 3ʹUTR-containing mRNAs in iPSC. C. steady-state read counts fold-change (y-axis) between long and short 3ʹUTR-containing mRNAs during neuronal development. D, mRNA half-life fold-change (y-axis) during neuronal development split according to distance (x-axis) between long and short 3ʹUTR-containing mRNAs. E, mRNA half-life fold-change (y-axis) separately for long and short 3ʹUTR-containing mRNAs split by developmental transitions. Genes with a distance >500 or <500 nucleotides between polyadenylation sites corresponding to long and short 3ʹUTR-containing mRNAs are included in the boxplots.

In agreement with published results (Mayr, 2016), we observed an accumulation of mRNAs with long 3ʹUTRs in NPC and neurons (Fig 5C). We next sought to quantify the contribution of mRNA stability to the apparent switch to long 3ʹUTR isoforms during neurodevelopment. We observed in iPSC that the half-life of mRNAs containing long 3ʹUTRs is less than in isoforms with short 3ʹUTRs when the difference in length is greater than 500 nt (Fig 5D). For example, a long 3ʹUTR isoform can be 4-fold less stable than its short isoform in iPSC (Fig 5D). Although less prominent in NPC and neurons, a similar trend of less stable long isoforms was observed in those cell types (Fig 5D).

As previously described (Rodrigues *et al*., 2016; Zheng *et al*., 2018), we expected that an increase in the half-life of the long 3ʹUTR isoforms in NPC and neurons would explain the increased abundance of these mRNAs relative to the short isoforms. Surprisingly, we observed a gradual destabilization of the short 3ʹUTR-containing mRNAs after both transitions (Fig 5E). Long 3ʹUTR-containing mRNAs decreased their half-life only after the transition from NPC-to-Neu (Fig 5E). Overall, the reduced half-life of the short 3ʹUTR isoforms was observed in the first transition to NPC but is most evident as an approximately 2-fold decrease in mRNA stability relative to the long 3ʹUTR isoforms when comparing iPSC-to-Neu. This effect on the short isoforms was observed when the difference in length between 3ʹUTR isoforms was > 500 nt but not when it was < 500 nt (Fig 5E). These data suggest that mRNA half-life has a differential impact on a subset of short compared to long 3ʹUTR-containing mRNAs during neurodevelopment.

### MicroRNA accumulation in neurons and destabilization of a subset of mRNAs with short 3ʹUTRs

To probe the contribution of microRNAs to the regulation of 3ʹUTR isoforms at a greater depth, we performed small RNA sequencing spiked with small exogenous RNA normalized by cell numbers to quantify relative and absolute fold-changes in microRNA abundance. Reassuringly, microRNAs specific to pluripotent stem cells (miR-302, miR-200) (Judson *et al*., 2009; Gill *et al*., 2011) and neurons (miR-137, miR-7) (Zahr *et al*., 2019) accumulated in the appropriate cell types after both transitions (Fig 6A, Supplementary table 6). Small RNA-seq data from the same control neuron samples have already been reported in comparison to isogenic RTT neurons (Rodrigues *et al*., 2023). As expected, we observed a remodeling of the microRNA landscape with 676 (25%) and 585 (22%) microRNAs changing steady-state abundance after iPSC-to-NPC and NPC-to-Neu transitions, respectively (Fig 6A). Interestingly, coincident with the shutdown of pluripotent stem cell-specific microRNAs, the number of microRNAs with a large fold-change (>5) is 7 times higher after the first transition from iPSC-to-NPC (14%) compared with NPC-to-Neu (2.2%).

**Figure 6.**
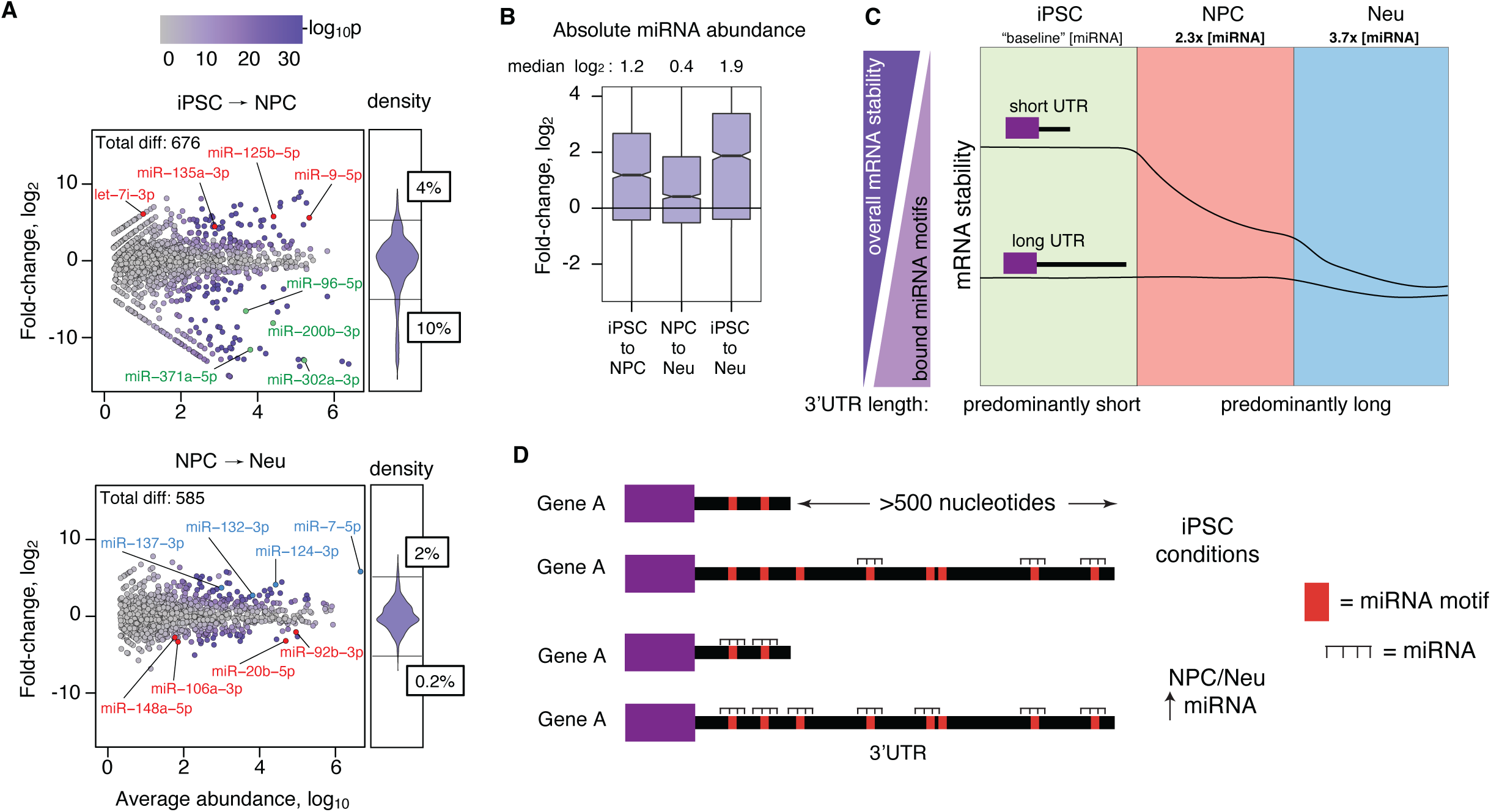
microRNA accumulation co-occurs with the destabilization of short 3ʹUTRs in cortical neurons. A, scatter plot showing changes in microRNA abundance during both developmental transitions. The x-axis represents the basal abundance of each mature microRNA detected, and y-axis represents their fold-change. Cell-type marker microRNAs for iPSC, NPC and neurons are highlighted in green, red and blue colours, respectively. B, DESeq2 microRNA (miRNA) steady-state level fold-change after spike-in normalization. Total RNA was extracted from the same number of cells, to which we added the same mass of a commercially available small RNA spike-in mixture. This spike-in mixture contains 54 different small RNA molecules covering a wide range of sequence compositions and molar concentrations. The absolute abundance of miRNAs increased for most miRNAs during neuronal development. C, summary of changes in short and long 3’UTR isoform stability during neurodevelopment. Gradients show mRNA stability in comparison to increased miRNA motifs present in the long isoform. D, model showing low miRNA abundance in iPSC favouring degradation of a subset of long isoforms, in contrast to higher miRNA abundance in NPC and neurons leading to instability of both isoforms.

As the microRNA profile changes during neurodevelopment, the half-life of their mRNA targets is expected to change. Indeed, the half-lives of mRNAs defined in TargetScan (Agarwal *et al*., 2015) as targeted by pluripotent stem cell-specific microRNAs like miR-302 and miR-200 were the most increased upon transition to NPC (Fig S6A). Further, normalization of microRNA levels using the small RNA spike-ins showed a ∼4-fold global increase in microRNA abundance per cell from iPSC-to-Neu, with the main increase during the transition to NPC (Fig 6B and Fig S6B). The increased microRNA abundance in NPC relative to iPSC was not accompanied by a change in mean half-life. However, the highest microRNA abundance in neurons (Fig 6B) was accompanied by the observed reduction of global mRNA half-life (Fig 2A).

The observed stability changes across neurodevelopment of the subset of short and long isoforms that differ by >500 nt are summarized in Figure 6C. The long isoforms already have a short half-life in iPSC and NPC, which declines modestly in neurons. In contrast, the short isoforms were more stable in iPSC and their half-life was reduced in NPC and neurons. This reduction in half-life co-occurred with the overall increased microRNA abundance per cell in NPC and neurons. The cumulative effect is that isoforms are predominantly short in iPSC and longer in neurons.

Taking the concept of microRNA thresholds into account, long isoforms will have more microRNA motifs than short isoforms (Fig 6D). When microRNA abundance per cell is low in iPSC, the long isoforms will be more susceptible to degradation than short isoforms. As the microRNA abundance per cell increases in NPC and neurons, there will be a minimal effect on long isoform degradation. In contrast, the increased microRNA threshold in NPC will have more substantial effects on short isoforms and reduce their half-life, until maximal degradation of short isoforms is established in neurons (Fig 6D).

### MicroRNA motifs enriched in buffered and boosted gene sets in NPC

We previously found no association of microRNA motifs with buffering in RTT neurons compared to control neurons (Rodrigues *et al*., 2023). The findings that per-cell microRNA levels increased during neurodevelopment prompted us to perform microRNA motif analyses on the most abundant isoform of the buffered and boosted gene sets. In iPSC-to-NPC, microRNA motifs were primarily enriched in the buffered genes with HL down and depleted on the buffered genes with HL up (Fig 7A, Supplementary table 4). In iPSC-to-Neu, only the microRNA motif depletion was observed in the buffered HL up genes (Fig 7A), and negligible microRNA motifs were enriched in NPC-to-Neu (Fig S4). This suggests that microRNAs were mostly deployed to reduce the half-life of buffered genes in the first transition, and that NPC and neurons only share a depletion of microRNA motifs on the buffered genes with HL up.

**Figure 7.**
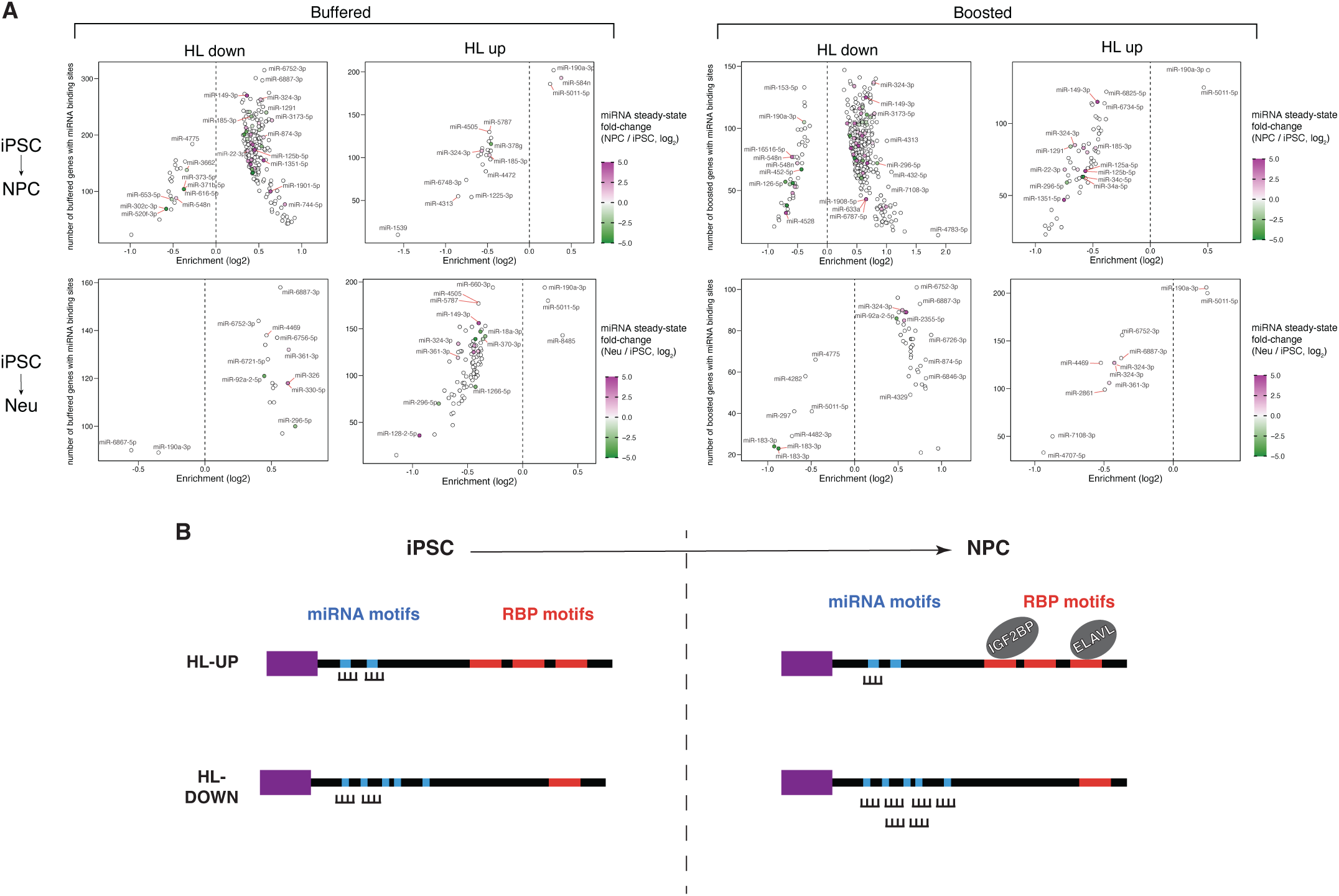
microRNA motifs are enriched on buffered and boosted genes during neurodevelopment. A, TargetScan miRNA 7-mer seeds enriched in the groups of buffered and boosted genes in iPSC-to-NPC (top) and iPSC-to-Neu (bottom) transitions. Colors represent the miRNA steady-state fold-change in the same neurodevelopmental transition shown in Fig 6A. B, model summarizing enriched and depleted miRNA and RBP motifs found on buffered and boosted genes, including the impact of miRNA abundance changes on the most abundant isoform in the iPSC-to-NPC transition. On half-life up genes (HL Up), miRNA sites are relatively depleted in NPC which have more miRNA abundance, and RBP motifs for stabilizing factors are enriched. On half-life down (HL-Down) genes, miRNA sites are relatively enriched in NPC and the stabilizing RBP motifs are depleted.

Strikingly, for boosted genes in iPSC-to-NPC, motifs for numerous microRNAs were also enriched or depleted in boosted genes with HL down, and some were depleted on boosted genes with HL up (Fig 7A, Supplementary table 5). Only a hint of enriched microRNA motifs were found in the iPSC-to-Neu comparison suggesting that this mechanism in NPC is maintained to a limited degree in neurons (Fig S4). Only two motifs were enriched on the boosted genes with HL up. Overall, these results generally implicate microRNAs in destabilizing the half-life of boosted and buffered genes during the iPSC-to-NPC transition, and this mechanism is partially maintained in the NPC-to-Neu transition. These microRNA motifs likely work in concert with the RBP motifs identified earlier leading to transcriptional boosting and buffering in human neurodevelopment (Fig 7B).

## DISCUSSION

We studied mRNA stability regulation to identify post-transcriptional mechanisms leading to transcriptional boosting and buffering during human neurodevelopment. We used RATE-seq to calculate relative transcript half-life changes as the ratio of standard RNA-seq steady-state and transcription rate measured using short 4sU incorporation times (Muhar *et al*., 2018). We found that roughly 20% of differentially regulated genes during transitions of iPSC-to-NPC, and NPC-to-neurons, were regulated only by changes in transcription rate. Strikingly, roughly 80% of differentially regulated genes experienced shifts in mRNA half-life. Half-life regulation was most commonly used to buffer transcription rate changes, leading to no significant change in steady-state levels, with approximately half of the genes in this set. The remaining genes were regulated either through half-life only or by boosting transcription changes. Boosted and buffered genes during human in vitro neurodevelopment were cross-validated using cell fractionation datasets (Blair *et al*., 2017).

These findings demonstrate that mRNA stability is an equal partner with transcription rate in establishing the steady-state levels of the transcriptome in human neurodevelopment. Given the consistently high frequency of buffered, boosted and half-life only gene sets in iPSC, NPC and neurons, it is highly likely that equivalent modes of gene regulation will be found when differentiating iPSC into other human cell types to model development and disease. Importantly, the same modes of gene regulation should be widely found in vivo, as we have already verified buffering and other gene regulation modes in a high-confidence RNA-seq resource of control and *Mecp2* null mouse brain samples (Boxer *et al*., 2020). We propose that during typical neurodevelopment, buffering reduces noise to increase robustness of gene expression while boosting accelerates gene upregulation or downregulation during differentiation.

We found evidence that similar RNA stability mechanisms cooperate with the core pluripotency transcription factors and adjust the abundance of their targets. In agreement with our data, a recent study showed that knockdown of *OCT4* or *NANOG* alters the expression of many RBP genes (Dvir *et al*., 2021). In addition, we detected buffering of roughly half of the transcriptionally upregulated targets. Overall, our data indicate the role of mRNA stability in buffering genes controlled by pluripotency transcription factors. Strikingly, only 30-40% of their target genes were regulated by transcription rate only. Our datasets may prove useful to define the transcription rate changes on target genes of other epigenetic or transcription factors implicated in human neurodevelopment and its disorders.

### MicroRNA abundance increases during neurodevelopment

We focused on the role of microRNAs because they can be quantitatively assessed by RNA-seq on a per-cell basis, and observed a global 4-fold increase per cell in cortical neurons compared to iPSC. Average mRNA half-life analysis revealed a global genome-wide reduction by roughly 2-fold in neurons compared to iPSC and NPC. We propose that increasing microRNA abundance on a per cell basis contributes to the mRNA degradation of most genes in neurons. We also detected genome-wide degradation of long 3’UTR-isoforms that have >500 nt more sequence than their corresponding short isoforms in pluripotent stem cells. This observation supports and expands the previously reported findings of the *MECP2* gene (Rodrigues *et al*., 2016). Our results suggest that microRNAs degrade a subset of long 3’UTR isoforms in pluripotent stem cells.

### Short isoform instability in NPC contributes to 3’UTR lengthening

Upon differentiation into NPC, we observed a striking destabilization of the short 3’UTR isoforms that differ by >500 nt from their long isoforms, that led ultimately to equivalent stability between long and short 3’UTR isoforms in neurons. Data from mouse hippocampal neurons (Tushev *et al*., 2018) corroborates the negligible difference in stability between long and short 3’UTR isoforms at this stage. A switch to more distal 3’-end cleavage sites through alternative polyadenylation participates in the overall increase in 3’UTR length consistently observed during neurodevelopment of different species (Hilgers *et al*., 2011; Smibert *et al*., 2012; Ulitsky *et al*., 2012). We confirmed the relative accumulation of long 3’UTR isoforms in neurons leading to an increase in the average 3’UTR length in these cells. However, we probed the accumulation of long 3’UTR isoforms at a greater temporal resolution than previously studied by including the measurements at the NPC stage. We detected a substantial contribution of mRNA instability to average 3’UTR length through the destabilization of a subset of short 3’UTR isoforms upon differentiation into NPC and neurons. Further, the considerable increase in relative long 3’UTR isoform abundances upon differentiation into NPC was again driven by an mRNA instability mechanism. Overall, our data shows that mRNA instability shifts contribute to the apparent lengthening of a subset of 3’UTRs in neurodevelopment that has its biggest effect at the NPC stage with an incremental increase in neurons.

### MicroRNA threshold effects may preferentially degrade short isoforms

Destabilization of short isoforms may involve a microRNA threshold (Mukherji *et al*., 2011; Ebert and Sharp, 2012). In iPSC, short and long 3’UTR-containing mRNAs are equally transcribed and processed. However, the long isoforms can be bound by many microRNAs (Fig 6D) that exceed the threshold level for degradation and could act as a competitive sponge for the few microRNAs that interact with short isoforms. Neuronal differentiation leads to an accumulation of microRNAs, shifting the threshold of available microRNAs per mRNA molecule (Bosson *et al*., 2014). This would lead to less sponging by the long isoforms that are still maximally degraded, resulting in increased degradation of the short 3’UTR-isoforms in NPC and maximal degradation in neurons. The result would be an increase in the average 3’UTR length, and it may also contribute to the decreased global mRNA half-life in neurons. Such stability-dependent regulation in pluripotent cells agrees with widespread *AGO2* binding to the transcriptome and the functional significance of microRNA biogenesis machinery on proper self-renewal and differentiation (Kanellopoulou *et al*., 2005; Wang *et al*., 2007; Leung *et al*., 2011; Li *et al*., 2020). Potential roles for RBPs in this process cannot be excluded in part because they can help disengage microRNA:target mRNA interactions (Kundu *et al*., 2012), but calculating global RBP abundance changes during differentiation at the level of absolute protein molecule number per cell will require exquisitely sensitive proteomic methods.

### Enriched motifs that stabilize mRNA on buffered and boosted genes in NPC and neurons

We previously associated RBPs that shuttle from the nucleus to the cytoplasm in buffering HL down genes in RTT neurons, but found no enriched motifs on buffered genes with HL up (Rodrigues *et al*., 2023). During neurodevelopment however, we now find enriched motifs for shuttling RBPs including ELAVLs, IGF2BPs, and PABPCs in both buffered HL up and boosted HL up genes. Similar RBP motif enrichments were found in buffered and boosted genes in the iPSC-to-Neu transition. Contrary to the RBP motifs, microRNA motifs were generally more enriched on the buffered and boosted HL down genes. These observations suggest a switch in post-transcriptional regulation by RBPs and microRNAs at the NPC stage (Fig 7B), that is maintained and further amplified by microRNA abundance changes in neurons.

### Limitations

A limitation to our approach is that we cannot detect changes in mRNA modifications and polyadenylation tail length that have been associated with buffering in cell lines (Slobodin *et al*., 2020). For example, N6 methylAdenosine (m6A) modifications are known to increase over neurodevelopment and are most abundant in neurons. m6A in 3’UTRs could serve to increase the number of targets for m6A-binding RBPs such as ELAVLs and IGF2BPs to increase half-life or influence 3’UTR length in neurons (Hilgers *et al*., 2012; Ke *et al*., 2015; Huang *et al*., 2018; Wei *et al*., 2020). A limitation to the motif analysis is that binding motifs are not known for all RBPs. It is the combinatorial and cumulative action of RBPs and microRNAs, taken together with additional yet-to-be-identified amplifier systems in neurons, that determine the mRNA half-life shifts regulating the levels of boosted and buffered transcripts and 3’UTR isoforms during neurodevelopment.

## Acknowledgments

This study was funded by grants from the Canadian Institutes of Health Research (CIHR; PJT-148746, PJT-168905, and ERARE Team ERT 161303 to J.E.); the Canada First Research Excellence Fund (Medicine by Design Cycle I: J.E.); the Col. Harland Sanders Rett Syndrome Research Fund at the University of Toronto (J.E.); the Ontario Brain Institute (POND Network: J.E.); and John Evans Leaders Fund/Canada Foundation for Innovation (JELF/CFI: J.E.); Canada Research Chairs Program (M.D.W. and J.E.); Beta Sigma Phi International Endowment Fund (J.E.); Early Researcher Award from the Ontario Ministry of Research and Innovation (M.D.W.); Genome Canada Disruptive Innovation in Genomics Grant (M.D.W. to support K.E.Y.); NIH grant R35GM128680 and the University of Colorado RNA Bioscience Initiative (O.R.); David Steven Cant Scholarship (M.M.); BioTalent Canada and Lunenfeld Summer Student awards (G.B.).

## Author contributions

Conceptualization, M.M., D.C.R., G.B., A.N., O.S.R., and J.E.; Investigation, M.M., D.C.R., G.B., A.N., K.Y., W.W., A.P., and J.L.; Software, M.M., G.B. and A.N.; Writing – original draft, M.M., D.C.R., G.B. and J.E.; Writing - Review & Editing, M.M., D.C.R., G.B., A.N., K.Y., P.P., O.S.R., B.T., M.D.W., and J.E.; Visualization, M.M., D.C.R. and G.B.; Supervision & Funding Acquisition, O.S.R., B.T., M.D.W., and J.E.

## Data availability

The original sequencing data that support the findings of this study can be accessed from GEO using the access number GSE191168 (wild type Neu), and wild type iPSC and NPC will be made available on publication. The hESC-derived data re-analyzed in this study can be downloaded from GSE100007 (nuclear and cytoplasmic subcellular fraction RNAseq).

## Code availability

The code used in this manuscript is available at GitHub https://github.com/JellisLab/stabilome-neurodevo

## SUPPLEMENTARY FIGURES

**Figure S1. Quality control of the RATE-seq experiment.** A, 4sU incorporation kinetic curves of human mRNA normalized to fly spike-in RNA. B, representative agarose gels showing 4sU-labeled pulled-down RNA for each time-point in iPSC and NPC. The presence of human (*Hs*) and fly ribosomal RNAs are denoted by arrows. C, read counts of spike-in RNAs: unlabeled yeast relative to fly RNA for all time points in iPSC, NPC, Neu. Unlabeled yeast RNA spike-in was used as a negative pull-down control (background control). The absence of yeast RNA indicates that the streptavidin-biotin pull-down of 4sU-labeled RNAs had minimal contamination of unlabeled human RNAs but was readily detectable in the steady-state samples. D, 4sU does not cause changes in neuronal viability or cell numbers at the dose and times used in the experiment. E, Pearson’s correlation between replicates of the human RNA for each time-point. F, Pearson’s correlation of ERCC spike-in RNAs between replicates (relative), and spike-in concentration and sequencing measured abundance (absolute) used for all time points in iPSC, NPC, Neu. ERCC spike-in RNA was used as a control for library prep quality. High Pearson’s correlations indicate the high quality of the library samples.

**Figure S2. Half-life measured with 4sU saturation curve and ratio method.** A, percentage of genes with measured half-life depending on transcription rate. B, the accuracy of half-life shown against half-life magnitude. CI_50_ stands for 50% confidence interval. Black points denote genes with poorly fit saturation curves and are removed from analysis in panel C. C, comparison of half-life estimated with two methods for well-measured genes selected in panel B.

**Figure S3. Subcellular fractionation from hESC-derived day-14 and day-50 neurons both validate buffered and boosted genes during neurodevelopment.** Number and percentage of all genes with changes in the 4 modes of regulation during cell transitions: buffered, half-life (HL) only, boosted and transcription rate (TR) only from hESC-derived NPC to day-14 (left panel) and day-50 (middle panel) neurons. Fraction of genes exhibiting the same mode of change (right panel) in the NPC-to-day-14 and NPC-to-day-50 neurons for each of the 4 modes of regulation. Overlaps were calculated using a hypergeometric test.

**Figure S4. Few RBP and microRNA motifs are enriched in buffered and boosted genes in NPC-to-Neu.** 6-mers known to be targeted by RBPs (top) and 7-mers targeted by microRNAs (bottom) enriched in the groups of buffered and boosted mRNAs in the NPC-to-Neu transition.

**Figure S5. Summary of long and short 3ʹUTR isoform quantification.** A, fraction of transcription rate reads assigned to the highest or top 2 highest in abundance 3ʹUTR isoforms. B, number of genes (Y-axis) depending on distance (X-axis) between long and short 3ʹUTR-containing mRNAs. C, representative sequencing read peaks in the iPSC transcription rate (upper) and steady-state (lower) samples, corresponding to polyadenylation sites (arrowheads) in the 3ʹUTR as an example of genes with active degradation of long 3ʹUTR-containing mRNAs. Y-axis represents the number of sequencing reads.

**Figure S6. Impact of microRNA on mRNA half-life.** A, Fold-change of mRNA targets of pluripotent-specific microRNAs (miRNAs) relative to all genes in iPSC-to-NPC transition depending on miRNA fold-change magnitude. B, fold-change for endogenous miRNAs and small RNA spike-ins after developmental transitions.

## Methods

### iPSC cultures, NPC, and neuronal differentiation

iPSC line #37 (WT) was previously described (Cheung *et al*., 2011). This cell line was generated and cultured under the approval of the SickKids Research Ethics Board and the Canadian Institutes of Health Research Stem Cell Oversight Committee. Methods to culture and differentiate the cell types are exactly as described (Rodrigues *et al*., 2023). In brief, iPSC were cultured on BD hESC-qualified matrigel (BD) in mTeSR medium (STEMCELL Technologies). To generate Neural Progenitor Cells (NPC), iPSC were aggregated as Embryoid Bodies (EBs) and replated after 7 days for growth in N2 media + laminin (1 ml/ml). After 7 days, neural rosettes were manually picked and transferred to poly-L-ornithine + laminin-coated wells and picked again after another 7 days. NPC were grown as a monolayer and split every 5-7 days in NPC media containing DMEM/F12, N2, B27, 1x non-essential amino acid (NEAA), 2 mg/ml Heparin, 1 mg/ml laminin. To generate neurons, NPC were plated on poly-L-ornithine + laminin-coated plates and cultured for 3 weeks in neural differentiation medium (Neurobasal, N2, B27, 1 mg/ml laminin, 1x penicillin-streptomycin, 10ng/ml BDNF, 10ng/ml GDNF, 200 mM ascorbic acid, and 10 mM cAMP).

### 4sU metabolic labeling and RNA extractions

The RATESeq experiment was performed exactly as described (Rodrigues *et al*., 2023). In brief, culture media of all three cell types was replaced with media supplemented with 100 μM 4sU (Sigma-Aldrich) reconstituted in DMSO. Cells were incubated at 37°C, 5% CO_2_, and total RNA was harvested at 0.5, 1, 4, 8, and 24 hours after the addition of 4sU. Metabolic labeling was designed such that all time points were collected together. Cells were quickly washed twice in ice-cold PBS 1x and lysed on the plate by addition of 1 mL of Trizol (Thermo Fisher Scientific). Total RNA was extracted according to manufacturer instructions. The steady-state samples were prepared from a 5μg aliquot of the 24 h time-point to which we added 0.5 μg of both unlabeled *yeast* and 4sU-labeled S2 fly *s*pike-in RNAs. Cellular viability in the presence of 100 μM 4sU was monitored up to 24 h of treatment on parallel cultures by using Trypan blue staining and live/dead cell counting.

### Biotinylation and pull down of 4sU-labeled RNAs

Biotinylation and pull down of labelled RNA was performed exactly as described (Rodrigues *et al*., 2023). In brief, 50 µg of total RNA was mixed with 5 µg unlabeled *yeast* RNA and 5 µg 4sU-labeled S2 fly RNA. 4sU Biotinylation was performed by adding 120 µL of 2.5× citrate buffer (25 mM citrate, pH 4.5, 2.5 mL EDTA) and 60 µL of 1 mg/mL HPDP-biotin (ThermoFisher Scientific) to the RNA samples for each time point. The solution was incubated at 37°C for 2 h. Samples were extracted twice with acid phenol, and once with chloroform. RNA was precipitated, pelleted and resuspended in 1× wash buffer (10 mM Tris-HCl, pH 7.4, 50 mM NaCl, 1 mM EDTA). Biotinylated RNAs were purified using the µMACS Streptavidin microbeads system (Miltenyi Biotec). 50 µL Miltenyi beads per sample were pre-blocked, applied to microcolumns and washed 5x. Beads were demagnetized and eluted off the column, and columns placed back on the magnetic stand. A total of 200 µL beads were mixed with each sample of biotinylated RNA and rotated at room temperature for 20 min. Samples were loaded into the microcolumns, washed 3x with wash buffer A (10 mM Tris-HCl, pH 7.4, 6 M urea, 10 mM EDTA), and washed 3x with wash buffer B (10 mM Tris-HCl, pH 7.4, 1 M NaCl, 10 mM EDTA). Final RNA samples were eluted with 5 washes of 1x wash buffer supplemented with 0.1 M DTT, and the flow-through was collected. Purified RNA samples were precipitated at −20°C overnight. Samples were spun and resuspended in 20 µL water. RNA quality was assessed by running 3 µL of samples on a 1.5% agarose gel.

### cDNA synthesis and qRT-PCR

cDNAs were synthesized from the nuclear RNA fractions using SuperScript III reverse transcriptase (ThermoFisher) with random hexamer primers according to the manufacturer’s instructions. For qRT-PCR, we used SYBR Select PCR Master Mix (ThermoFisher). ΔCt values of the target genes were normalized against the spike-in RNAs from yeast and fly genes. Absolute fold-change measurements relative to cell numbers were calculated using the 2^−(ΔΔCt)^ method, averaged between technical and subsequently biological replicates to achieve an average fold difference. The primers used are described in the primers table.

### miRNA extraction and spike-in strategy

To calculate relative and absolute differences in the miRNA population in iPSC, NPC and Neurons, small RNAs were extracted from two replicates of both lines using the same number of cells followed by the addition of a set of spike-in RNAs exactly as described (Rodrigues *et al*., 2023). Small RNAs were extracted from 500,000 cells of each line using the SPLIT RNA extraction Kit (Lexogen). A set of 52 RNA spike-ins (QIAseq miRNA Library QC Spike-Ins – Qiagen) spanning a wide range of molar concentrations were added to the recovered RNAs. Sequencing libraries were made using the Small RNA library preparation kit NEBNext (NEB). Sequencing was performed on the Illumina HiSeq 2500 using the Rapid Run mode.

### Library preparation and RNA-sequencing

RNA-seq libraries were prepared for each time-point and steady-state sample exactly as described (Rodrigues *et al*., 2023). In brief, the QuantSeq 3ʹ mRNA-Seq Library Prep Kit FWD for Illumina (Lexogen) was automated on the NGS WorkStation (Agilent) at The Centre for Applied Genomics (TCAG). PCR cycle numbers were determined using the PCR Add-on Kit for Illumina (Lexogen). All steady-state samples were processed with 250 ng of total RNA input. To minimize variability between time-points within a batch, RNA samples were processed with the same total RNA input with a minimum of 100 ng of total RNA used. Each sample was spiked-in with ERCC RNA Spike-In Control Mix 1 (Ambion) prior to the start of library preparation. Library quality and quantity were measured at The Centre for Applied Genomics (TCAG) with Bioanalyzer (Agilent) and KAPA qPCR (Roche). Sequencing was performed at TCAG on the Illumina HiSeq 2500 with single-end 100bp read length yielding 40 to 50 million reads.

### Processing of raw sequencing reads

Processing was performed exactly as described (Rodrigues *et al*., 2023). In brief we trimmed reads in 4 steps using cutadapt version 1.10. First, we removed adapters exactly at the 3ʹ-end of the reads. Second, we removed internal or long stretches of adapter. Third, we trimmed low-quality bases at the 3ʹ-end of the reads. Finally, we removed poly-A tail at the 3ʹ-end of the reads.

### Generation of custom hybrid genome index and reads alignment with STAR

We generated a custom genome index to accommodate the quantification of yeast, fly, and ERCC spike-in RNA exactly as described (Rodrigues *et al*., 2023). Annotations (gencode version 29, flybase version all-r6.22, saccharomyces_cerevisiae.gff from yeastgenome.org, custom for ERCC) and genomes (hg38, dm6, sacCer3, ERCC from ThermoFisher) for all species and ERCC were combined and then processed with STAR version 2.6.0c (--sjdbOverhang 100). Finally, reads are aligned to hybrid genome with STAR version 2.6.0c (default settings) (Dobin *et al*., 2013).

### Quantification of RNA abundance

Poly-A sites were obtained from PolyA_DB version 3 and converted to hg38 coordinates with *liftOver* (UCSC) (Karolchik *et al*., 2004; Wang *et al*., 2018). Reads with MAPQ < 2 were filtered out. Finally, usage of poly-A sites was defined as a sum of reads whose 3ʹ-ends are falling within 20bp upstream and 10bp downstream of the poly-A sites. The sum was counted with a custom Python script using *pybedtools*, *pysam*, *pypiper*. (Dale *et al*., 2011; Danecek *et al*., 2021; Sheffield *et al*., 2021). The abundance of mature miRNAs was quantified with *mirdeep2* pipeline (Friedländer *et al*., 2012). Reads were preprocessed and collapsed with *mapper.pl* script (-e -h -j - k AGATCGGAAGAGCACA -l 18 -m -v) and quantified with *quantifier.pl* script, using hairpin and mature sequences obtained from miRbase (Kozomara *et al*., 2019).

### Normalization of human read counts with fly spike-ins

First, reads were assigned as originating from either human, fly, yeast or ERCC, based on alignment to hybrid genome. Then, human raw counts were divided by the sum of all fly spike-in raw counts. Since both human and fly RNAs are 4sU-labeled, this normalization to fly spike-ins reconstructs the fraction of 4sU-labeled human RNA at each time point.

### Spike-in RNA usage clarification

Fly 4sU spike-ins were used to normalize pulled down human counts for all time points (0.5-hour to 24-hour) to assist with transcription rate and absolute half-life calculations. Yeast spike-ins were used as quality control for contamination in the pull-downs. ERCC spike-ins were used to control for sequencing quality. First, it directly estimates sequencing error magnitude, excluding biological variation. Second, it shows the capacity of 3ʹ-end QuantSeq to reconstruct molar concentrations. Note that the steady-state sample was not normalized with spike-ins and is separate from the 24-hour time point sample.

### Transcription rate and half-life measurements

The transcription rate was estimated from 0.5-hour and 1-hour time points exactly as described (Rodrigues *et al*., 2023). This estimate assumes that RNA degradation is negligible for most genes at time points before 1 hour. First, human normalized counts at 1-hour time point are divided by 2 to create a new approximate replicate at 0.5-hour time point. Division by 2 is done to account for a twice longer period of transcription in a 1-hour time point sample. To clarify, the addition of an approximate replicate was motivated by the reduction in the standard error of log_2_FC estimated with *DESeq2*. To compare the transcription rate between cell types using *DESeq2*, human normalized counts are further quantile normalized between replicates of the same cell type with *normalize.quantiles* (*preprocessCore* package) (Love *et al*., 2014). Half-life was estimated in 2 separate ways: fit of the 4sU saturation curve and the ratio of steady-state to transcription rate. For the 4sU saturation curve method, the half-life is estimated in a 2-step procedure. First, normalized counts are fit with *nls* (nonlinear least squares from *stats* package) to approximate the true number of counts Y at each timepoint. Then, in a second pass, normalized counts are fit again with *nls*, but now correcting for the increase in variance using weights set as 1/Y. Confidence intervals are estimated with *confint* function (*stats* package). For the ratio method, the half-life is estimated with *DESeq2* using raw human counts from 0.5-hour, 1-hour and steady-state samples (design =∼ assay). The assay is a 2-level factor one for transcription rate and one for steady-state. 0.5-hour and 1-hour samples correspond to the transcription rate.

### The shift in average mRNA half-life between cell types

To describe the global absolute mRNA half-life shift, we filtered out unreliable genes from the saturation curve method. First, both gene transcription rate and steady-state should be above the bottom 10% quantile. Second, the ratio of 50% confidence interval and estimate of the half-life should be below 0.75. Then, the mean half-life was calculated from the filtered set of genes.

### Random forest prediction of up and down-regulated genes in transcription rates

Features of the classifier are frequencies of k-mers in 3ʹUTR or gene-body, calculated using *oligonucleotideFrequency* (*Biostrings* package). Predicted variable denotes genes that are either up or downregulated in transcription rate. Thresholds for the human data were:

1. TR_up_: log_2_FC_TR_ > 1 & padj < 0.1
2. TR_down_: log_2_FC_TR_ < −1 & padj < 0.1

Data was split into 75% and 25% for training and test sets. The classifier is trained using *randomForest* (*randomForest* package).

### Transcription factors targets data

Targets of transcription factors NANOG, OCT4, and a repressive complex PRC2 defined by ChIP-seq were downloaded from molecular signatures database (MSigDB) (Subramanian *et al*., 2005; Ben-Porath *et al*., 2008; Liberzon *et al*., 2011).

### Processing of cell culture datasets from Blair et al 2017 Cell Reports

Nuclear eardand cytoplasmic RNA-seq data for hESC, NPC, and neurons were downloaded from *GSE100007*. Half-life was estimated as a ratio between cytoplasmic counts and nuclear counts using the interaction term approach in *DESeq2* (design = ∼ celltype + batch + assay + celltype:assay) as described (Rodrigues *et al*., 2023). Overlaps between RATE-seq and subcellular fraction data (Blait *et al*., 2017) was calculated for each individual mode of regulation using a hypergeometric test, (phyper in R) using a background of genes expressed in either of the cell types being tested with TPM > 1.

### Transite analysis of miRNAs and RBPs

Genes were split into groups and multiple comparisons of 3’UTR sequences between groups were performed as described (Rodrigues *et al*., 2023) for each neurodevelopmental transition. Groups of compared genes:

1. Foreground: TRdown and HLup Background: TRdown
2. Foreground: TRup and HLdown Background: TRup
3. Foreground: TRdown and HLdown Background: TRdown
4. Foreground: TRup and HLup Background: TRup

### Computer code

All computer code used for sequencing analysis is available on GitHub at https://github.com/JellisLab/stabilome-neurodevo

### Primer list for qRT-PCRs

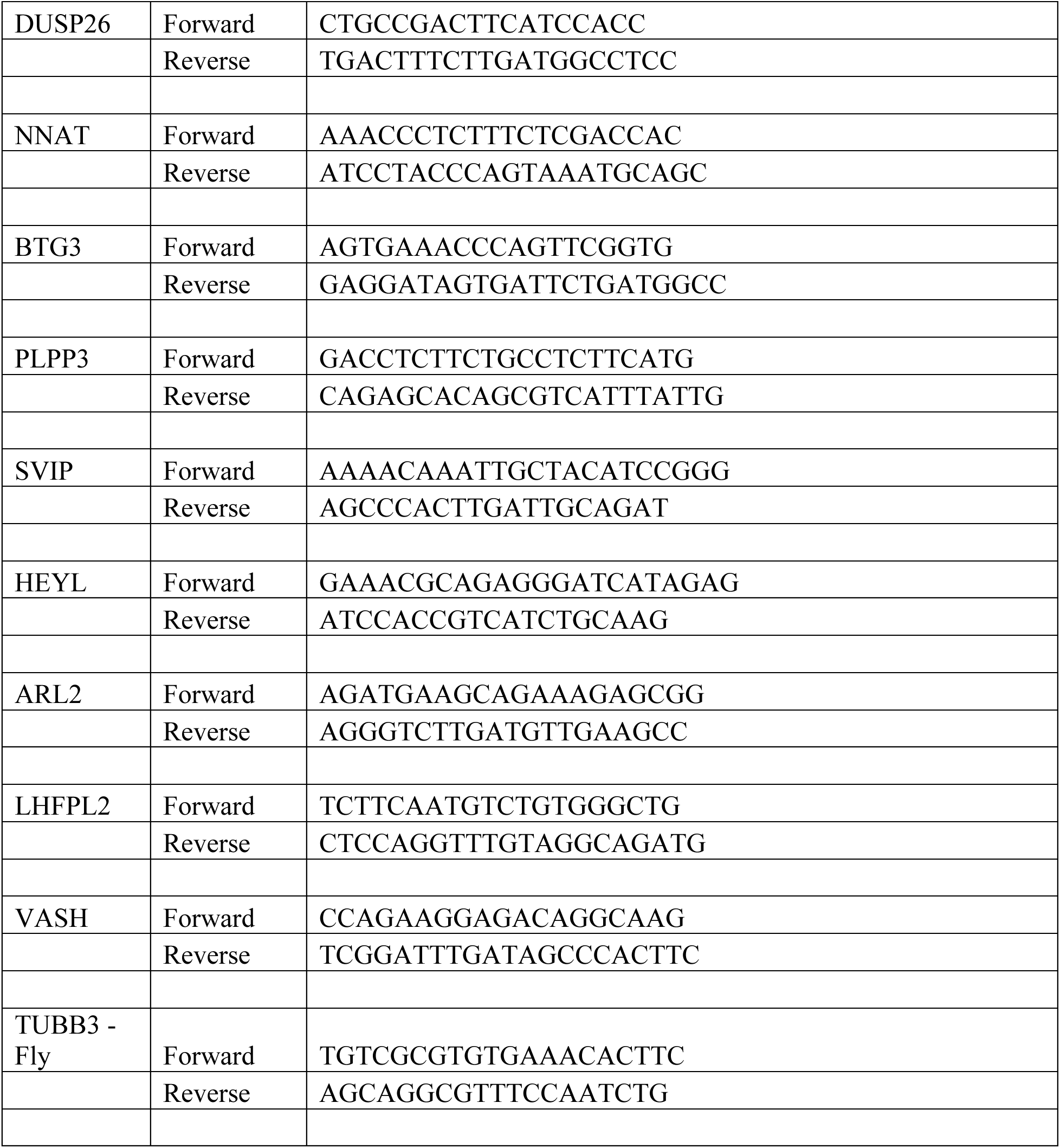
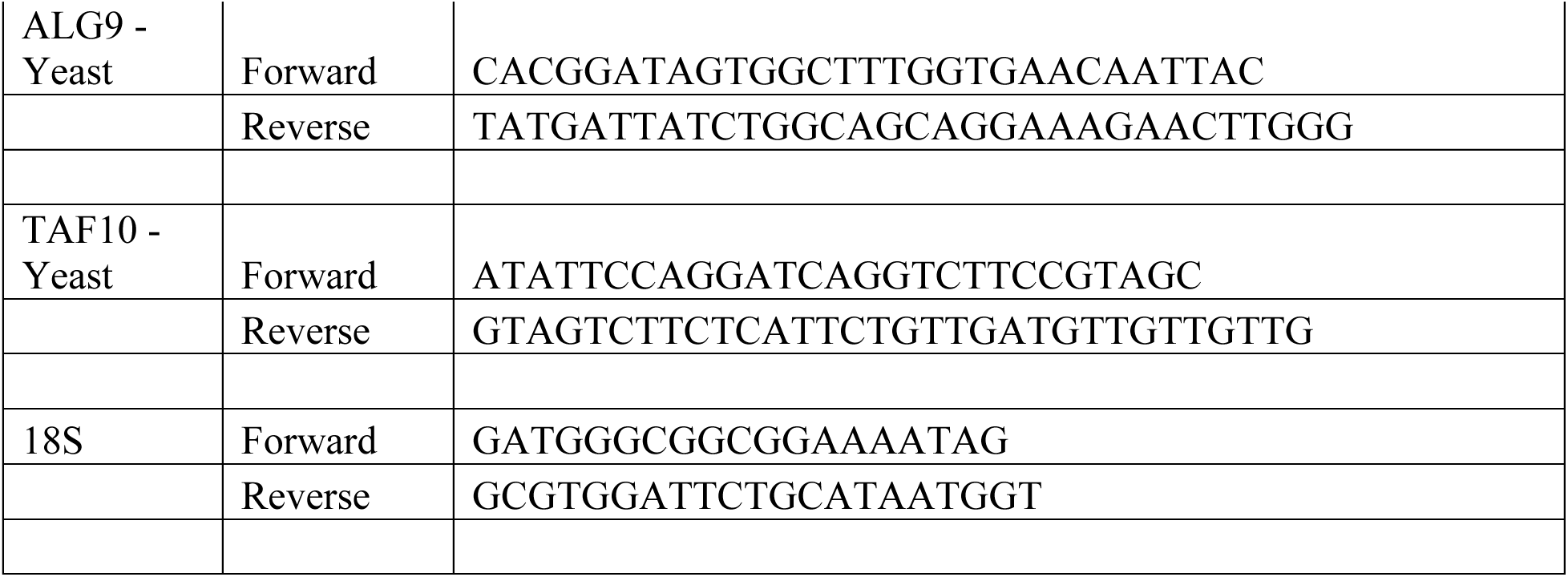

