## Supplementary figures and images for "Transcriptional boosting and buffering during neurodevelopment by shifts in mRNA instability"

### Supplementary Figures

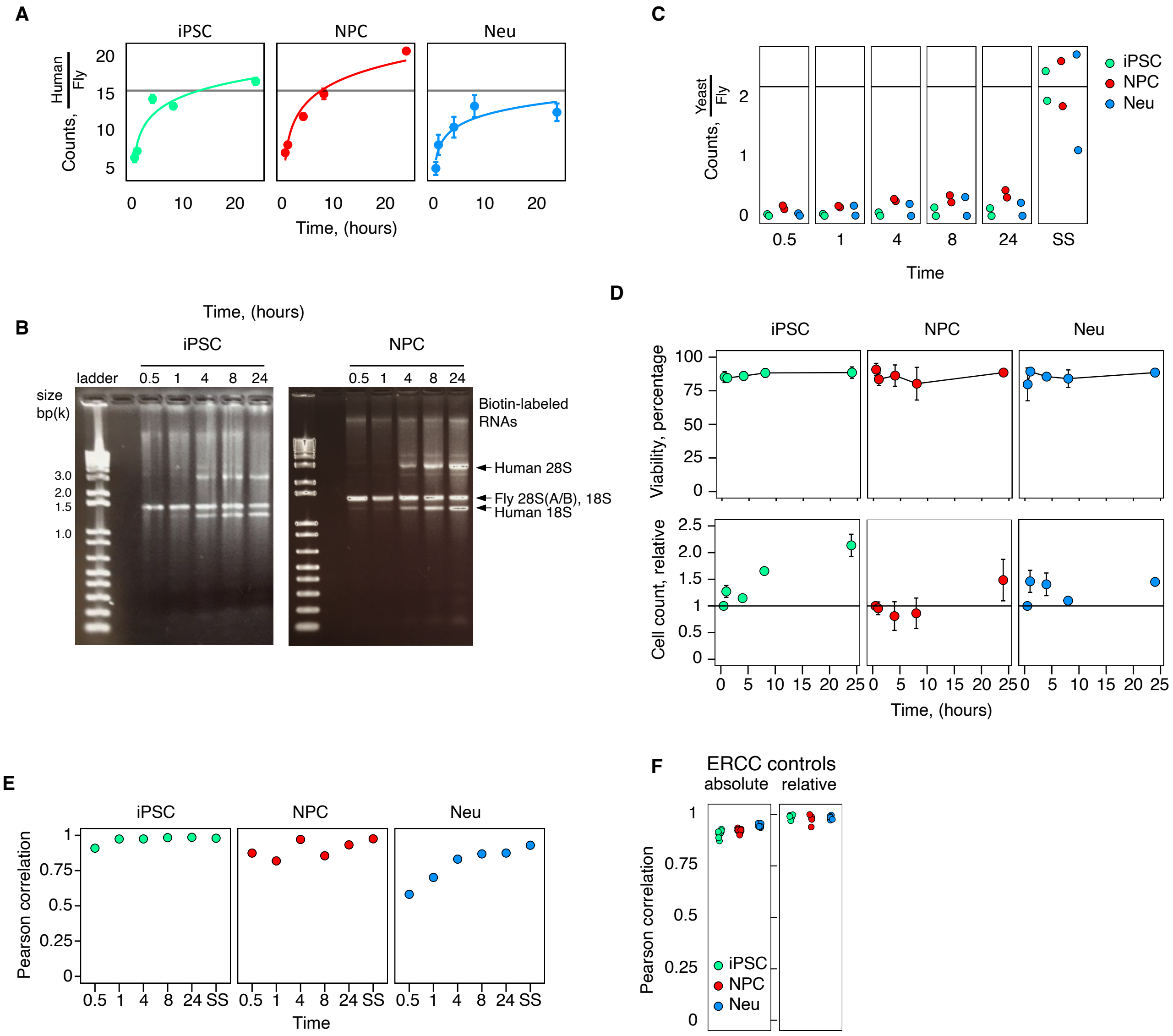

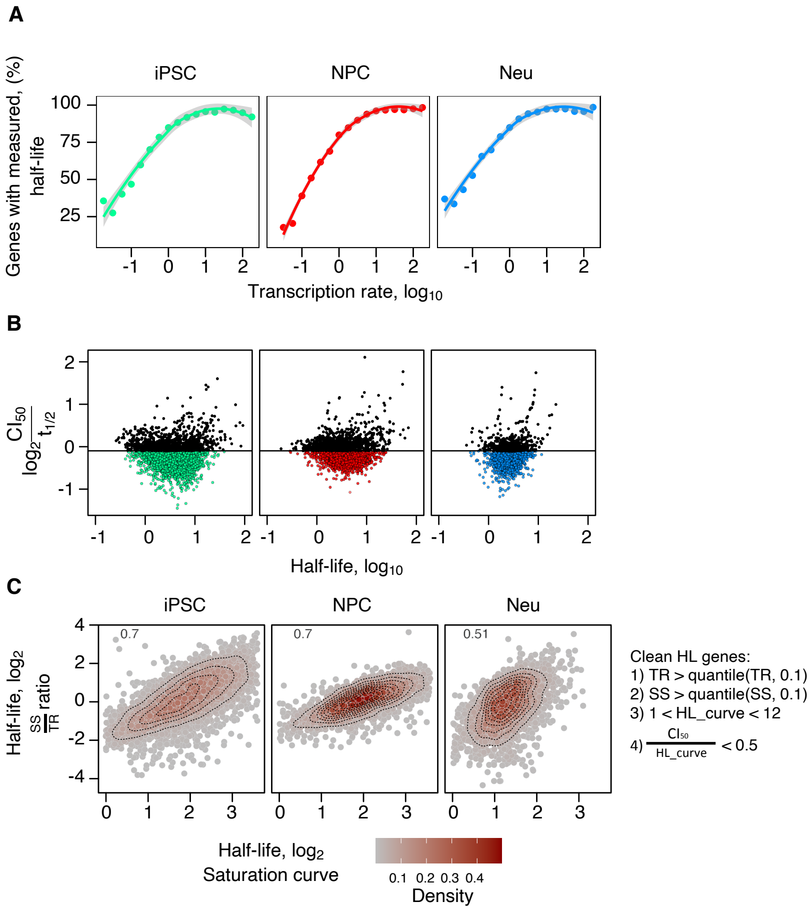

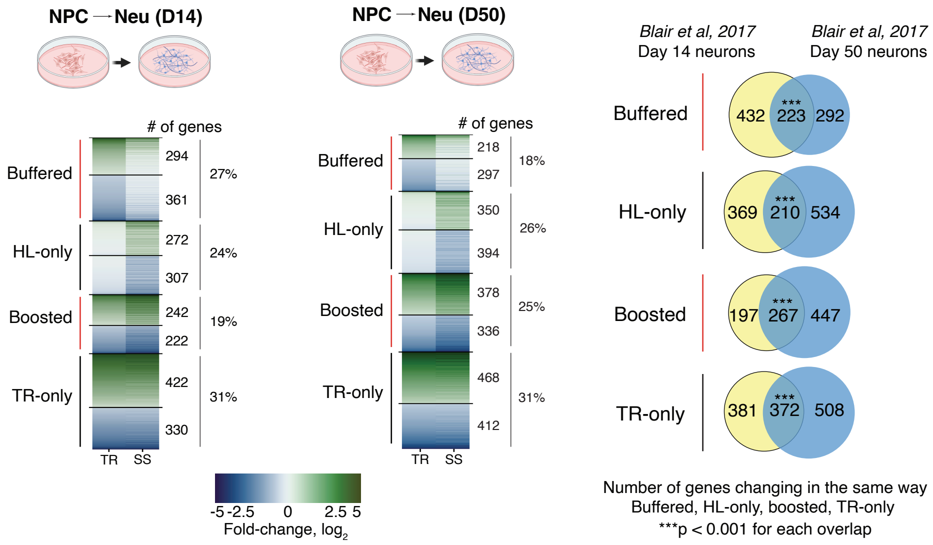

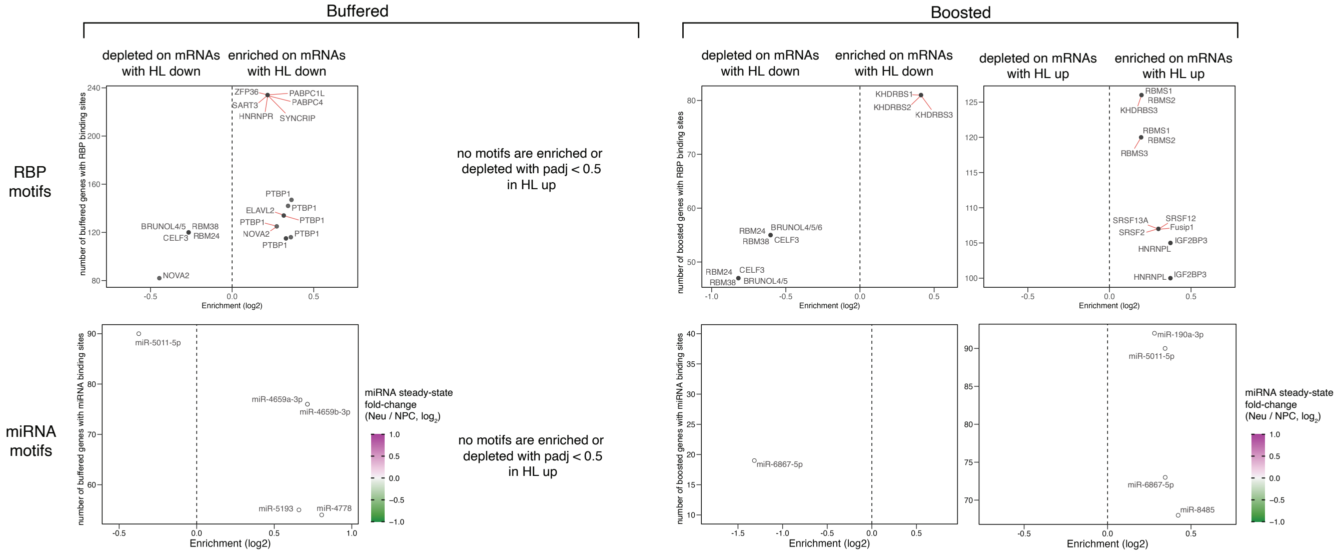

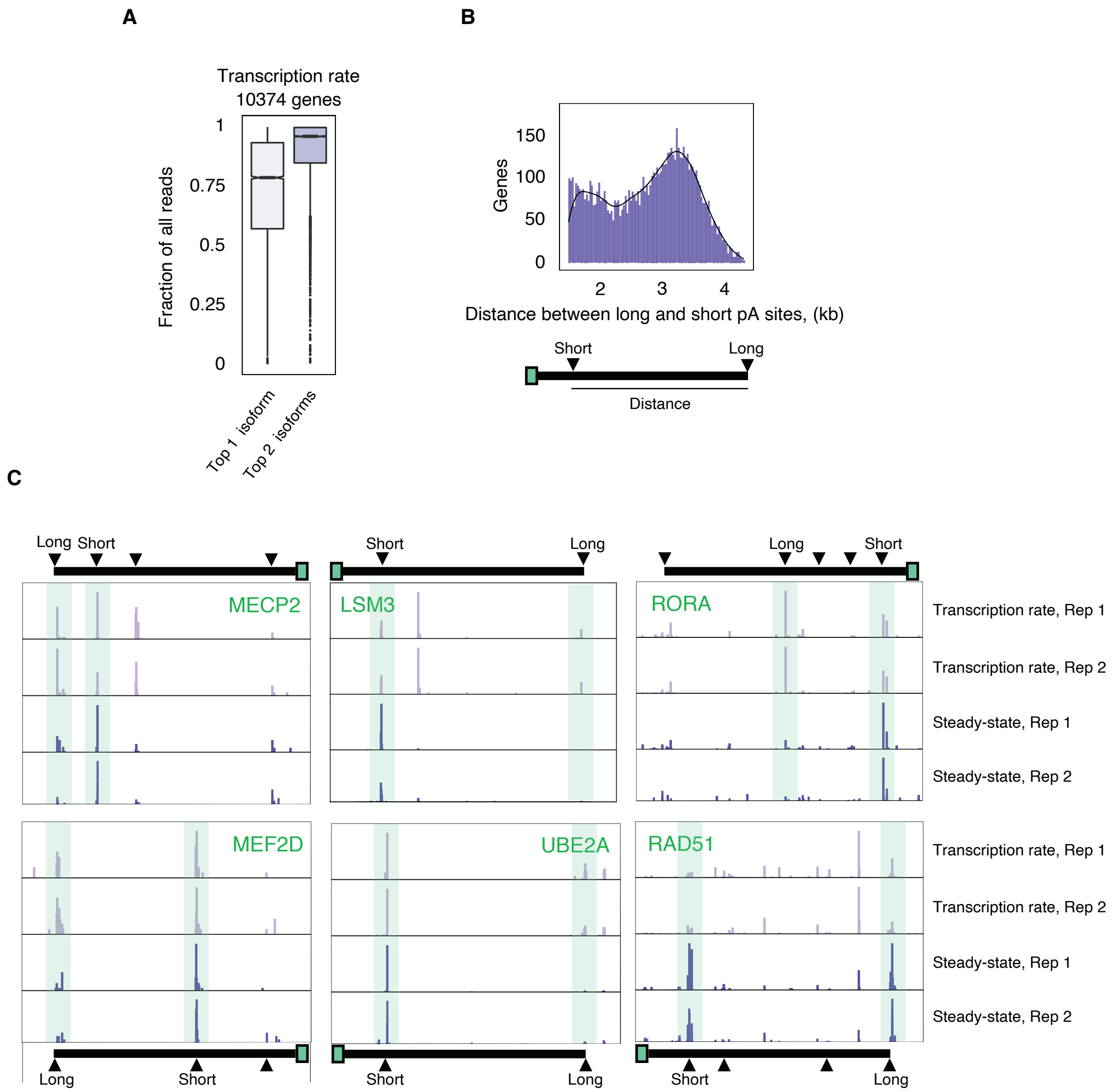

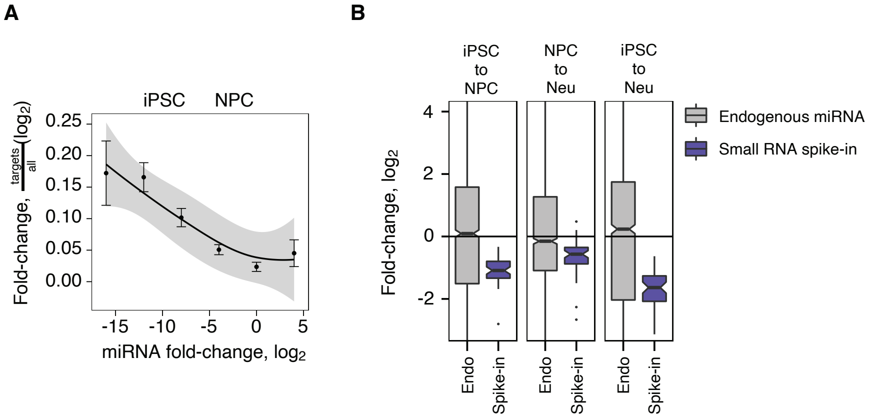
